# Executive function supports single-shot endowment of value to arbitrary transient goals

**DOI:** 10.1101/2020.10.21.348938

**Authors:** Samuel D. McDougle, Ian C. Ballard, Beth Baribault, Sonia J. Bishop, Anne G.E. Collins

## Abstract

People often learn from the outcomes of their actions, even when these outcomes do not involve material rewards or punishments. How does our brain provide this flexibility? We combined behavior, computational modeling, and functional neuroimaging to probe whether learning from transient goals harnesses the same circuitry that supports learning from secondary reinforcers. Behavior and neuroimaging revealed that “one-shot” transient goals (abstract fractal images seen once) can act as a substitute for rewards during instrumental learning, and produce reliable reward-like signals in dopaminergic reward circuits. Moreover, we found evidence that prefrontal correlates of executive control may play a role in shaping these responses in reward circuits. These results suggest that learning from abstract goal outcomes is supported by an interplay between high-level representations in prefrontal cortex and low-level responses in subcortical reward circuits. This interaction may allow humans to perform reinforcement learning over flexible, arbitrarily abstract reward functions.

## INTRODUCTION

During real-world goal-directed behavior, successful actions are often not signaled by immediate receipt of primary or secondary reinforcers (e.g. food, money) but through the realization of goals. Consider a child first learning to play basketball – the first successful shot through the hoop should, in theory, elicit a positive response and boost the value of the preceding action. In this example, the achievement of an abstract, novel goal (i.e., getting the ball through the hoop for the first time) could be translated into a conventional reinforcer. For that translation to occur upon her first ever success, her brain has to rapidly endow value to this abstract goal using a single experience, and her internal determination that this outcome is indeed a goal. This impressive aspect of human behavior – our ability to rapidly endow value to an abstract object or event – is, in our view, taken for granted. Indeed, this ability contrasts with even our closest primate cousins: Chimpanzees can learn to treat novel tokens as substitutes for reward, but they do so only after lengthy bouts of conditioning where those tokens are associated with food (Cowles, 1937; Wolfe, 1936). Rapid, one-shot endowment of arbitrary goals with value is likely a unique faculty of higher-level human cognition.

Parallels between learning from abstract novel goals versus traditional secondary reinforcers (e.g., extra pocket money for tidying your bedroom) are poorly understood. Here we test whether attaining novel, transient goals can reinforce choices in a similar manner to obtaining “points” (a commonly used secondary reinforcer), and examine the role of executive function in this process. We pushed this concept to its logical extreme, asking if goals can substitute for secondary reinforcers during learning even when those goals are abstract stimuli with no prior meaning or value to the learner, and are only observed a single time.

Our hypothesis is that standard reinforcement learning circuits support learning from arbitrary transient goals. This prediction is motivated by the well-known observation that there is a remarkable diversity of primary and secondary reinforcers that drive instrumental learning. Primary rewards, including intrinsically pleasant odors (Howard et al., 2015) and flavors (McClure et al., 2003), act as reliable reinforcers that engage striatal circuits. Secondary reinforcers, such as money or numeric “points” (Daw et al., 2006), which acquire value from repeated pairings with reward, also engage this system. More abstract secondary reinforcers, such as improvements in perceived social reputation (Izuma et al., 2008), words and symbols that explicitly signal outcome valence (Daniel & Pollmann, 2010; Hamann & Mao, 2002), and internally maintained representations about performance accuracy (Han et al., 2010; Satterthwaite et al., 2012), all consistently engage striatal learning systems. Lastly, information itself can act as a reward: When people attain outcomes that resolve uncertainty about the task they are performing, mesolimbic reward areas are activated (Charpentier et al., 2018; White et al., 2019). Taken together, decades of findings suggest that striatal populations operate according to a flexible definition of reward that is context-dependent, and includes arbitrarily abstract goals (Juechems & Summerfield, 2019).

Recent evidence suggests that sensitivity to goals can sometimes even supersede reward-related responses: Frömer and colleagues (2019) observed that responses in the brain’s value network are sensitive to achieving explicit, stable task goals, even when those goals do not involve – or are even in opposition to – rewarding outcomes (Frömer et al., 2019). It remains to be established how this system endows values to goals. Repeated experience with goal-relevant outcomes may engage incremental reward learning circuits to turn goal-related outcomes into secondary reinforcers, in the same way that social cues or numerical points acquire secondary value over time. Alternatively, the executive system may rapidly (i.e., in a single trial) imbue outcomes with value via top-down input. In order to adjudicate between these mechanisms, we designed a study in which both goals and goal-outcomes change from trial-to-trial.

Our hypothesis is also inspired by recent research demonstrating that top-down inputs directly influence value-based learning computations in RL circuits (Rmus et al., 2021). For instance, attention modulates RL by specifying reward-predicting features of stimuli that RL should operate on (Leong et al., 2017; Radulescu et al., 2019). Explicit memories of familiar rewards can be flexibly combined to endow value to novel combinations of those rewards, and to drive activity in the brain’s valuation network (Barron et al., 2013). Finally, reward prediction error responses in dopamine neurons are influenced by latent task representations that are likely maintained in the prefrontal cortex (Babayan et al., 2018; Schuck et al., 2016; Sharpe et al., 2019; Starkweather et al., 2018; Wilson et al., 2014), as well as information held in working memory (Collins, 2018; Collins et al., 2017; Collins & Frank, 2018). This body of work suggests that higher-level cognitive processes can specify and shape the key inputs (e.g., states, rewards, etc.) to low-level RL systems.

One line of research that also targets people’s ability to rapidly act on arbitrary goals is “Rapid Instructed Task Learning” (Cole et al., 2013). These tasks focus on the ease with which humans can leverage simple verbal (or symbolic) instructions to successfully perform novel tasks within a single trial. For instance, if a person is instructed to tap their left foot when they see a star-shaped line drawing, they will typically succeed at this task on the first try (Cole et al., 2013). Interestingly, damage to the lateral prefrontal cortex can disrupt this kind of behavior even when instructions are easily understood and remembered, producing so-called “goal neglect” (Duncan et al., 1995, 1996). PFC-mediated executive function appears to play a role in converting novel linguistic or symbolic inputs into “task sets” that can immediately guide action (Cole et al., 2013; Collins & Frank, 2013; Miller & Cohen, 2001). As mentioned above, it is unclear if learning from such rapidly formed task sets leverages conventional reward learning hardware or relies on its own dedicated circuit.

In this experiment, we used behavioral, computational modeling, and neuroimaging to investigate whether canonical reward signals measured during learning from familiar secondary reinforcers are observed, in overlapping neural regions, during learning from single-shot goal outcomes. We then examined whether prefrontal correlates of executive function, defined via a secondary executive function task, influence how reward learning regions respond to single-shot goal attainment.

## RESULTS

Subjects (N = 32) performed a probabilistic selection learning task (Frank et al., 2007) adapted for fMRI, and an executive function task (n-back; Kirchner, 1958). Each trial of the selection task required a binary choice between two stimuli (Figure 1A). Subjects were required to learn which stimulus in each pair more often produced favorable outcomes. The task involved two feedback conditions: Familiar and Single-shot. The Familiar condition used numeric points (i.e., “+1” or “+0”) as secondary reinforcers. Points, like money, are secondary reinforcers due to their common association with positive outcomes in various real-world games and activities; indeed, a multitude of RL studies have used points as proxies for reward, which lead to reliable learning behaviors and robust responses in the reward system (e.g., Daw et al., 2006). Therefore, we refer to the Familiar feedback condition as generating “reward outcomes.” The Single-shot feedback condition used randomly-generated fractal stimuli as outcomes (“goal” versus “non-goal”). To induce rapid learning from novel goals, and to prevent any particular goal stimulus from incrementally accruing instrumental value, both the goal and non-goal fractals were novel on each Single-shot condition trial. Thus, the Single-shot condition required subjects to use their knowledge of the current fractal goal to determine whether their choices were successful. Each trial of both conditions included three phases: a pre-choice phase, a choice phase, and a feedback phase (Figure 1A). In the Single-shot condition, the two novel fractal stimuli were displayed during the pre-choice phase with one designated as the goal and the other as the non-goal, explicitly mirroring, respectively, the “+1” and “+0” of the Familiar condition. To match motivational aspects of the two conditions, no performance-based monetary incentives were provided in the experiment.

**Figure 1.**
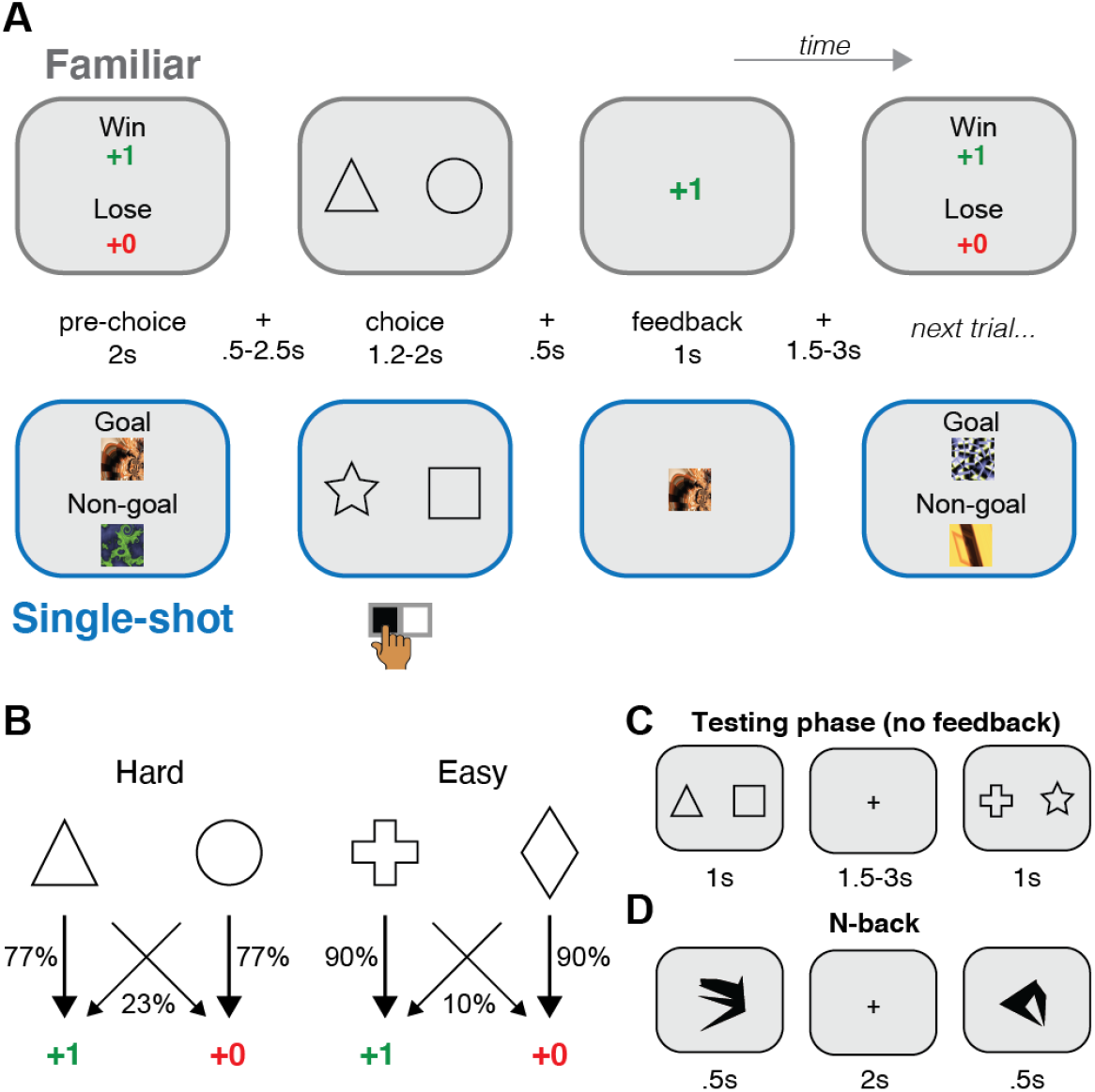
Task design. **(A)** Subjects (N = 32) performed a probabilistic selection task, learning which of two choice stimuli (e.g., triangle versus circle) was more likely to yield positive feedback. Two feedback conditions were used: in the Familiar reward condition, successful choices were rewarded with numeric points. In the Single-shot goal condition, successful choices were signaled via pre-specified “goal” fractal images. The images used as goal and non-goal stimuli were unique for each Single-shot trial. Pairs of choice stimuli were assigned to a single condition, and trials from each condition were intermixed. **(B)** In both feedback conditions, each choice pair was designated as either Hard or Easy, determined by the difference in success probabilities between pairs of choice stimuli. **(C)** After the learning phase of the probabilistic selection task, subjects experienced a testing phase, where different pairings of the 8 total choice stimuli were pitted against one another and subjects rapidly chose their preference. No feedback was given in this phase. **(D)** Subjects also performed an N-back task, in which they responded when an image within a sequence repeated after “N” intervening images. N’s used = [1,2,3].

To investigate differential effects of task difficulty and feedback condition, we also implemented two difficulty levels (Hard and Easy), where each pair of choice stimuli was associated with different probabilities of yielding successful outcomes (Figure 1B). To test subjects’ retention of learned values, a subsequent testing phase was administered where subjects made choices between each possible pairing of the eight learning stimuli and no feedback was given (Figure 1C). Finally, to capture an independent measure of executive function, subjects also performed a standard visual n-back task (Figure 1D; *N*s used: 1, 2, 3). We predicted that performance on this task would be specifically related to subjects’ ability to learn from novel goals.

### Executive function supports single-shot reward learning

Subjects performed well in the learning task, selecting the better stimulus of each pair in both the Familiar (mean: 82%; chance = 50%; *t*(30) = 19.39, *p* < 1e-17) and Single-shot (mean: 75%; *t*(30) = 10.70, *p* < 1e-11) conditions (Figure 2A, B). A repeated-measures ANOVA revealed a main effect of feedback condition (i.e., Single-shot versus Familiar; *F*(1,30) = 11.67, *p* = 0.002), with better learning phase performance in the Familiar versus the Single-shot condition (Bonferroni-corrected; *t*(30) = 3.41, *p* = 0.002). We also observed a main effect of difficulty (*F*(1,30) = 22.74, *p* < 1e-4), with better performance on Easy versus Hard trials (*t*(30) = 4.77, *p* < 1e-4). There was no significant interaction between feedback condition and difficulty (*F*(1,30) = 0.34, *p* = 0.57). These results show that subjects could leverage one-shot goal stimuli to successfully learn to select actions that lead to goals, but that this was less successful than learning from familiar rewards.

**Figure 2.**
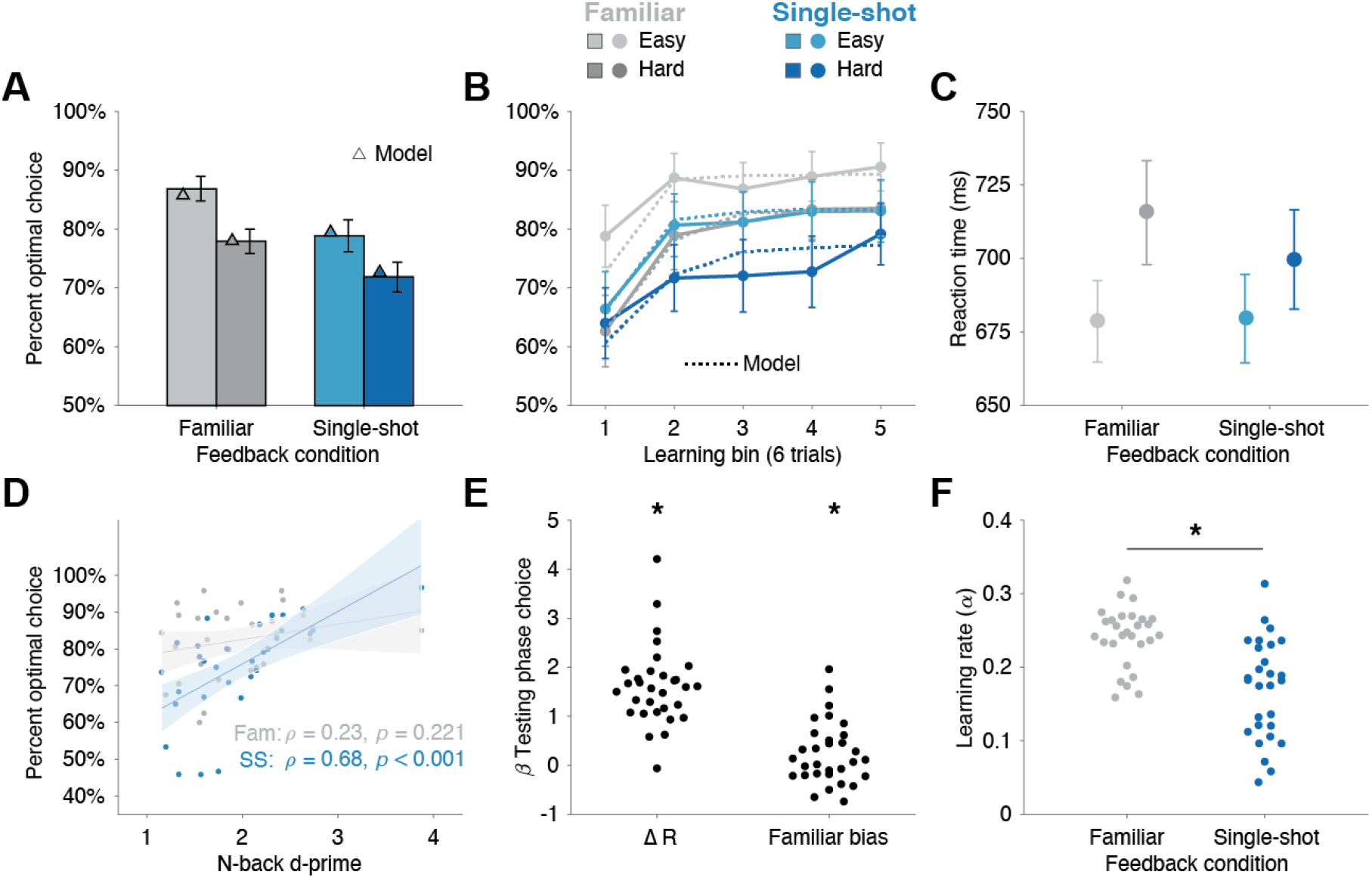
Behavioral results. **(A)** Average performance in the probabilistic selection task for each condition. Subjects performed well above chance (50%) in the task. We observed main effects of feedback condition (Familiar vs Single-shot) and difficulty (Easy versus Hard) but no interaction. Model simulation results (triangles) are depicted for each condition and difficulty level. **(B)** Learning curves for each condition and difficulty level, with 6 trials per bin. Model simulations are depicted as dashed lines. **(C)** Mean reaction times during learning. We observed a main effect of difficulty, but no effect of feedback condition nor any interaction. **(D)** Correlation of performance in the N-back task with learning performance in the probabilistic selection task (collapsed across difficulty levels). Fam = Familiar condition; Inst = Single-shot condition. **(E)** Regression weights resulting from a logistic regression analysis on testing phase choices. Subjects’ choice of stimuli increased as a function of how often the stimulus was rewarded during learning (ΔR, left), and showed a bias toward stimuli associated with Familiar reward feedback when they were paired with stimuli associated with Single-shot goal feedback (right). **(F)** Learning rate parameters from the best-fit RL model. Learning rates were significantly higher in the Familiar condition. * = *p* < 0.05; Error shading = 95% C.I.; Error bars = 1 s.e.m.

One explanation for performance differences between the Familiar and Single-shot feedback conditions is the dual-task demands in the latter (i.e., holding the novel “goal” fractal in memory while also having to select a preferred stimulus). A common signature of dual-tasks is slowed reaction times (RTs) for individual sub-tasks (Pashler, 1994). Thus, one prediction of a dual-task effect is slower RTs during choice in the Single-shot condition relative to the Familiar condition. An ANOVA on the RT data (Figure 2C), however, revealed no main effect of feedback condition (*F*(1,30) = 0.57, *p* = 0.47), a main effect of difficulty (*F*(1,30) = 14.07, *p* = 0.001), and no significant interaction (*F*(1,30) = 1.83, *p* = 0.19). These results show that the dual-task design of the Single-shot reward condition did not manifest by slowing RTs, which suggests that maintaining a novel goal image did not necessarily interfere with choice. These results also argue against a qualitatively different process (such as planning) driving behavior in the Single-shot condition: In general, goal-directed reasoning should incur an RT cost relative to simple instrumental learning (Keramati et al., 2011).

We hypothesized that the primary factor differentiating performance between the conditions was the fidelity of working memory. That is, if the goal fractal stimulus is sufficiently encoded and maintained in memory during the Single-shot trials, it will effectively stand in as a reward. If this is true, we expect working memory performance to correlate with Single-shot condition performance above and beyond performance in the Familiar condition (Figure 2D). Indeed, Single-shot condition performance was significantly correlated with n-back d-prime values (*ρ* = 0.68, *p* < 1e-4) but Familiar condition performance was not (*ρ* = 0.23, *p* = 0.221). A permutation test revealed that these correlations were significantly different (*p* = 0.038, 5000 iterations of shuffled correlation differences). We also found a significant correlation between n-back performance and the learning difference between feedback conditions (i.e., Single-shot minus Familiar; *ρ* = 0.52, *p* = 0.003). Nonparametric (Spearman) correlations were used for the above correlations given the one clear n-back task outlier (Figure 2D); however, we note that the results were replicated with parametric (Pearson) correlation metrics.

We next asked if executive function covaried with learning in the Single-shot condition simply because it was a harder (i.e., dual) task, or if the particular memory demands of the Single-shot condition required executive function. That is, the correlation results (Figure 2D) could arise due to simple differences in the difficulty of the learning task between conditions, as measured by choice performance. We controlled for difficulty by selecting difficulty-matched subsets of data – specifically, we examined Hard trials of the Familiar condition and Easy trials of the Single-shot condition, where performance was statistically indistinguishable (Bayes Factor = 6.84 in favor of the null). If n-back performance covaries with Single-shot condition performance for reasons beyond simple task difficulty, the correlation results in Figure 2D should hold in these data. Indeed, Single-shot-Easy performance was significantly correlated with n-back performance (*ρ* = 0.74, *p* = 0.004) but Familiar-Hard performance was not (*ρ* = 0.06, *p* = 0.504), and these correlations were significantly different (permutation test: *p* = 0.046). Taken together, these results suggest that executive function played a selective role in maintaining transient goals and helping subjects learn from them.

### Learning from single-shot goals versus familiar rewards is similar, but slower

How well were learned stimulus values retained after training was complete? One potential consequence of learning purely via top-down executive function – a plausible hypothesis for the Single-shot condition – is relatively brittle value representations that are forgotten quickly (Collins, 2018). On the other hand, if learning proceeds similarly between the conditions, the amount of forgetting should be the same. We addressed this question in the testing phase (Figure 1C). When looking at pairs of testing phase stimuli that were learned under the same feedback condition, we found that subjects selected the more valuable stimulus more often for both the Familiar and Single-shot stimuli (mean: 68%; *t*(30) = 8.49, *p* < 1e-8; mean: 64%; *t*(30) = 5.04, *p* < 1e-3; respectively), with no significant difference between feedback conditions (*t*(30) = 1.11, *p* = 0.28).

To characterize choice behavior across the full range of testing phase trial types, we further analyzed subjects’ choices using multiple logistic regression (see *Methods*). The choice of stimulus in the testing phase was influenced by the difference in the cumulative number of successful outcomes associated with each stimulus (‘ΔR’ in Figure 2E; *t*-test on Betas relative to zero: *t*(30) = 11.17, *p* < 1e-11), but we did not observe a significant interaction between cumulative value and the effect of the feedback condition in which the stimuli were learned (*t*(30) = 0.21, *p* = 0.839). This suggests that subjects similarly integrated both types of outcomes (rewards and goals) into longer-term memories of stimulus value. Lastly, when the two stimuli in the testing phase had originally been learned via different feedback conditions, subjects did show a bias toward stimuli from the Familiar condition, even when controlling for cumulative reward differences within the same regression (Figure 2E; *t*(30) = 2.16, *p* = 0.039). This suggests that values learned via familiar rewards may have been more salient during recall when directly pitted against those learned via novel goals.

Next we asked if performance differences between feedback conditions of the learning task resulted from choice- or learning-related effects. In order to better understand the differences between the feedback conditions, and to produce RL model regressors for fMRI analysis, we modeled subjects’ choices in this task with variants of standard RL models (Sutton & Barto, 1998). We implemented a Bayesian model selection technique (Piray et al., 2019) that simultaneously fits and compares multiple candidate models (see *Methods*). This analysis strongly favored a simple RL model that differentiated the Familiar and Single-shot conditions via separate learning rate parameters (Exceedance probability for this model versus competing variants = 1; Figure S1; Table S1). Critically, this model outperformed a competing model that used different levels of decision noise in each feedback condition – this suggests that the condition differences we observed (Figure 2A, B) were related to learning rather than choice, the latter being a natural prediction of dual-task interference.

To validate our model we simulated choices using the fit model parameters: As shown in Figure 2A and 2B, the model successfully reproduced subjects’ performance across feedback conditions and difficulty levels. Performance differences were successfully captured by the learning rate parameter – learning rates were significantly lower in the Single-shot condition versus the Familiar condition (*p* < 1e-4 via an “HBI *t*-test”, see *Methods*; Figure 2F and Figure S1). Moreover, consistent with the results depicted in Figure 2D, n-back performance was positively correlated with the difference between the Single-shot and Familiar condition learning rates (i.e., Single-shot minus Familiar; *ρ* = 0.44, *p* = 0.019).

We note that the observed difference in learning rates could represent (at least) two non-mutually-exclusive phenomena: First, it could be that there are weaker appetitive signals for novel goal stimuli versus familiar rewards. Second, occasional “lapses’’ in working memory could lead to forgetting of the fractal stimuli. The fact that n-back performance was selectively predictive of the Single-shot condition performance appears to support the lapsing interpretation, though executive function could also act to boost the appetitive strength of goals. Either way, choice and RT analyses suggest qualitatively similar, though slower, learning from single-shot goals versus familiar rewards. Next, we asked if these similarities carried over to the neural signatures of learning.

### Similar neural regions support familiar reward and single-shot goal-based learning

We reasoned that one-shot valuation of novel goals leveraged the same RL circuits that drive learning from familiar rewards, and that activity in executive function-related regions of the prefrontal cortex (PFC) could support this process through an interaction with these reward-sensitive regions. These results would be consistent with our behavioral results, where executive function performance covaried with goal-based learning (Figure 2D).

We first used whole-brain contrasts to measure univariate effects of feedback condition. In the pre-choice phase, we observed significantly more activity in the Single-shot versus Familiar condition in areas across the ventral visual stream, hippocampus, and both medial and lateral regions of PFC (Figure S2; see also for results of Familiar > Single-shot contrasts). These results are broadly consistent with both the greater visual complexity in the Single-shot condition during the pre-choice phase, where text-based instructions and a complex fractal stimulus are viewed, rather than simply text alone (Figure 1A). Additionally, there were increased cognitive demands during this phase in the Single-shot condition – subjects needed to attend to and encode the novel fractals. In the choice phase, we observed more activity in the medial striatum and visual cortex in the Instruced versus Familiar condition. The lack of any significant differences in PFC activation during the choice phase in this contrast is consistent with the relatively modest working memory demand in the Single-shot feedback condition (Figure S2). However, we note that the continued activation in primary visual areas in the Single-shot feedback condition during the choice phase could potentially reflect ongoing working memory maintenance (Emrich et al., 2013). Finally, in the feedback phase, we observed greater activation in the visual cortex and dorsolateral prefrontal cortex in the Single-shot versus Familiar condition. These increased activations could reflect, respectively, the complex visual features of the fractal stimulus and recall of its valence (i.e., goal or non-goal; Manoach et al., 2003).

We used ROI analyses to test the hypothesis that overlapping neural populations encode value signals related to traditional secondary reinforcers and novel goals. We used the Familiar feedback condition as a reward processing localizer task, generating the ROIs we used to characterize goal processing in the Single-shot feedback condition. Thus, our analysis of the Single-shot condition was fully validated out-of-set (see *Methods*). This approach provided a stringent test of our hypothesis that overlapping neural populations would encode reward and value signals related to both traditional secondary reinforcers and novel, transient goals. Moreover, ROIs for individual subjects were determined using a leave-one-out procedure where the group functional map used to create a particular subject’s functional ROI excluded that subject’s own data (Boorman et al., 2013; Kriegeskorte et al., 2009; see *Methods* for further details). Figure 3A shows example ROIs derived from the Familiar condition for one representative subject.

**Figure 3.**
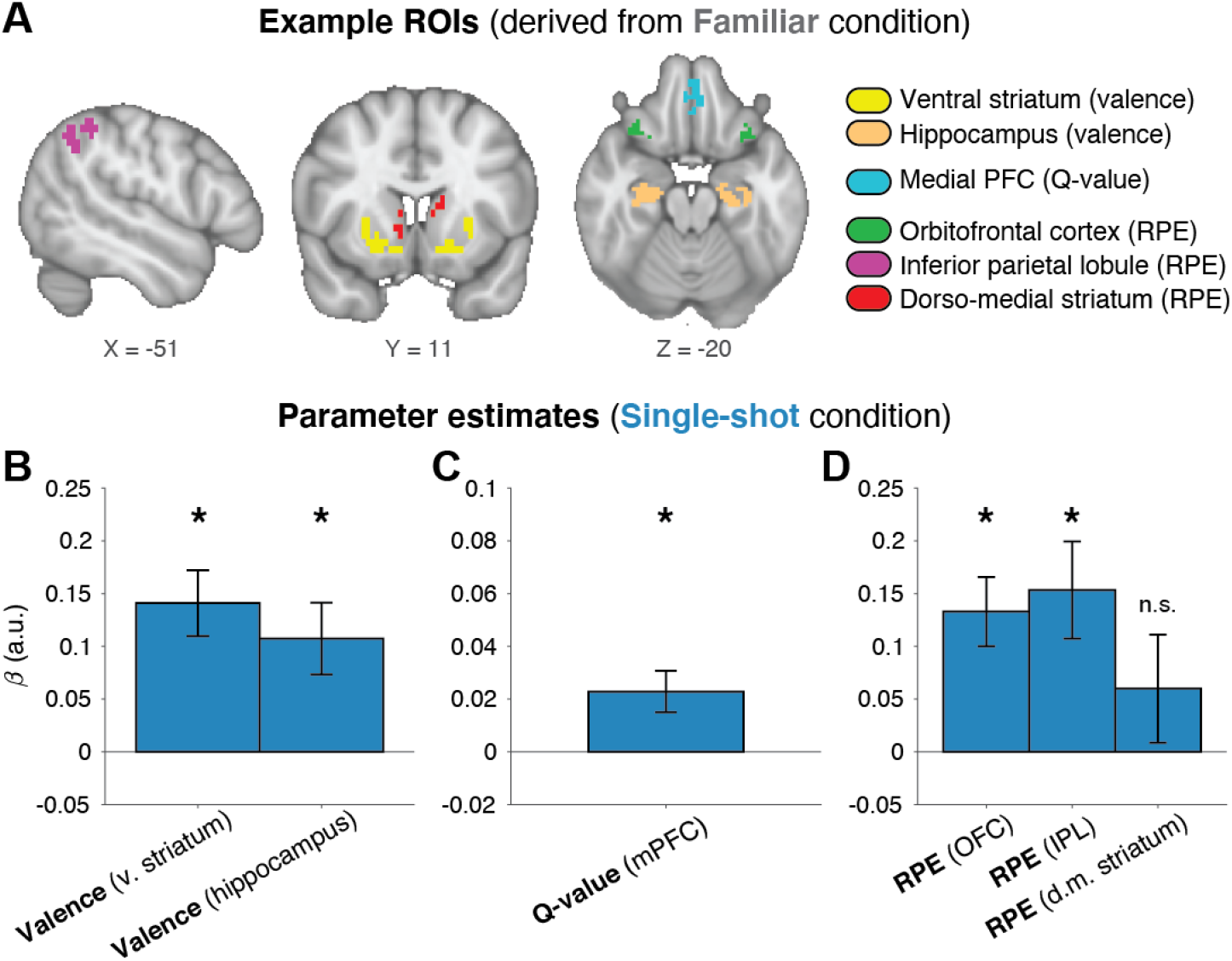
fMRI results: Valence, value, and prediction errors. **(A)** Example ROIs for the three main effects of interest are shown for an individual subject (subject s020). ROIs were created using a leave-one-out procedure, where each subject’s data were excluded from the statistical maps used to define their ROIs. Critically, only trials from the Familiar condition were used to generate these ROIs. Held-out Single-shot condition results were then computed in these ROIs, for effects of **(B)** valence, **(C)** model-derived Q-values at choice, and **(D)** model-derived reward prediction errors (RPEs). All parameter estimates were significantly greater than zero at *p* < 0.01, except where noted. Error bars = 1 s.e.m. mPFC = medial prefrontal cortex; OFC = orbito-frontal cortex; IPL = inferior parietal lobule.

We first sought to test whether subcortical valence responses were comparable across feedback conditions. In the Familiar condition, we observed predicted effects of feedback valence in the anterior hippocampus (HC; *t*-test on neural beta values: *t*(27) = 5.78, *p* < 1e-5) and the ventral striatum (VS; *t*(27) = 4.20, *p* < 1e-3), in addition to various cortical activations (Figure S2). Both of these subcortical results are consistent with previous findings of reward processing in RL-related circuits (Davidow et al., 2016; Delgado et al., 2000; Li et al., 2011; McClure et al., 2004; Palombo et al., 2019). Crucially, in the held-out Single-shot condition, valence responses were observed in both the hippocampal (*t*(27) = 3.16, *p* = 0.004) and VS (*t*(27) = 4.51, *p* < 1e-3) ROIs defined from the Familiar condition valence response (Figure 3B). Moreover, whole-brain contrasts revealed no clusters (cortical or subcortical) with significant differences in valence reponses between conditions (Figure S2). These results suggest that rapid endowment of value to a novel, transient goal stimulus leads to valenced responses in the same regions of the brain that show sensitivity to conventional reinforcers.

We hypothesized that if the brain’s reinforcement system is harnessed to learn from novel transient goals, the same computational latent variables should be present in both feedback conditions. The value of the selected choice stimulus (termed “Q-values” in standard RL models) typically correlates with activation in the medial frontal cortex (Bartra et al., 2013). In the Familiar condition, we observed the predicted Q-value coding in a ventral-rostral region of medial prefrontal cortex (mPFC; *t*(27) = 4.26, *p* < 1e-3; Figure 3A). Consistent with our predictions, we observed significant Q-value coding in this same mPFC ROI in the held-out Single-shot condition (Figure 3B; *t*(27) = 2.90, *p* = 0.007). This result implies that comparable computational processes are involved in modifying value representations through both familiar rewards and novel goals, and using those values to make decisions.

Reward prediction errors (RPEs) drive reinforcement learning, and have robust neural correlates in multiple brain regions. In the Familiar condition, we observed significant RPE-linked activity in various regions (Figure 3A, Figure S2), notably in a frontal ROI that included regions of bilateral orbitofrontal cortex (OFC, with some overlap in the insula; *t*(27) = 5.49, *p* < 1e-5), a second cortical ROI spanning bilateral portions of the inferior parietal lobe (IPL; *t*(27) = 4.98, *p* < 1e-4), and a bilateral ROI in dorso-medial striatum (DMS; *t*(27) = 3.99, *p* < 1e-3). These ROIs are consistent with findings from previous studies modeling the neural correlates of prediction errors (Daw et al., 2011; Garrison et al., 2013).

We observed significant effects in the two cortical RPE ROIs localized in the Familiar condition when analyzing the held-out Single-shot condition (Figure 3B; SPL: *t*(30) = 3.33, *p* = 0.003; OFC: *t*(30) = 4.06, *p* < 1e-3), further supporting the idea that comparable computational mechanisms drove learning in both feedback conditions. Moreover, whole-brain contrasts revealed no significant differences in RPE processing between the conditions (Figure S2). However, contrary to our prediction, the RPE response in the dorso-medial striatum ROI, while numerically positive on average, were not significantly greater than zero in the Single-shot condition (Figure 3B; *t*(30) = 1.17, *p* = 0.253). Control analyses using slightly different striatal ROIs also failed to reach significance (Figure S3). We note that this analysis of RPE-related activity is particularly conservative because the RPE regressor is included in the same model as – and thus competes for variance with – the outcome valence regressor. Consequently, significant activity in this analysis must reflect parametric RPE encoding beyond the effect of outcome valence.

To further probe if striatal RPEs were detectable in the Single-shot condition, we opted to take a cross-validated encoding-focused approach (see *Methods* for details). If the computations underlying RPEs in response to familiar rewards are mirrored during learning from goals, we should be able to decode goal RPEs in the striatum using a model trained on the Familiar condition data. We extracted feedback-locked betas for each individual trial of the learning task (from voxels in the striatal RPE ROI), and restricted our analysis to rewarded trials only. For each Familiar condition run, we trained linear models separately for each voxel, computing the ability of the feedback response amplitude to explain variance in the model-derived RPEs. RPEs were then decoded from the held-out BOLD data for both Single-shot and Familiar Condition runs, and compared to the associated held-out model-derived RPEs those same runs.

Cross-validated RPE encoding was observed within-condition across-runs (*t*(27) = 3.31, *p* = 0.003), between-condition within-run (*t*(27) = 6.39, *p* < 1e-5), and between-condition across-runs (*t*(27) = 2.34, *p* = 0.027). We emphasize that the regression models used for the encoding analyses were trained on Familiar runs only, providing a stringent test for RPE encoding in the Single-shot condition. These results suggest that goal prediction errors are represented in the same format as typical reward prediction errors in the dorso-medial striatum. However, we caution that these results were not mirrored in the conventional GLM analysis (Figure 3B). This discrepancy suggests that goal RPE signals in the DMS may be relatively weaker (or noisier) than familiar reinforcer RPE signals, consistent with our observation of slower learning in the Single-shot condition. However, we do note that Single-shot condition RPE signals in the frontal and parietal ROIs were statistically robust (Figure 3B).

### Single-shot goal learning drives increased frontal-striatal and frontal-hippocampal functional correlations

Our second hypothesis was that executive prefrontal cortex (PFC) regions encode and maintain novel transient goals, and interact with reward circuits so that those goals can act as reinforcers. We predicted that PFC activity during the encoding of goals would covary with reward system activity during feedback. In addition, we also predicted that this relationship would be stronger during goal-driven versus reward-driven learning. We specified an executive function prefrontal ROI by extracting working memory load-sensitive clusters from an independent n-back task (Figure S4). Load-sensitive clusters spanned regions of the precentral gyrus, middle frontal gyrus, and inferior frontal gyrus (Yeo et al., 2015). To isolate executive areas involved in goal processing in the learning task, we crossed the n-back ROI with a leave-one-out functional ROI of goal-encoding (Single-shot > Familiar contrast on the pre-choice phase, Figure S2; see Figure 5A for an example PFC ROI). For subcortical areas, we used the three ROIs defined previously (Figure 3A; VS and hippocampal valence-based ROIs, and the DMS RPE-based ROI).

**Figure 4.**
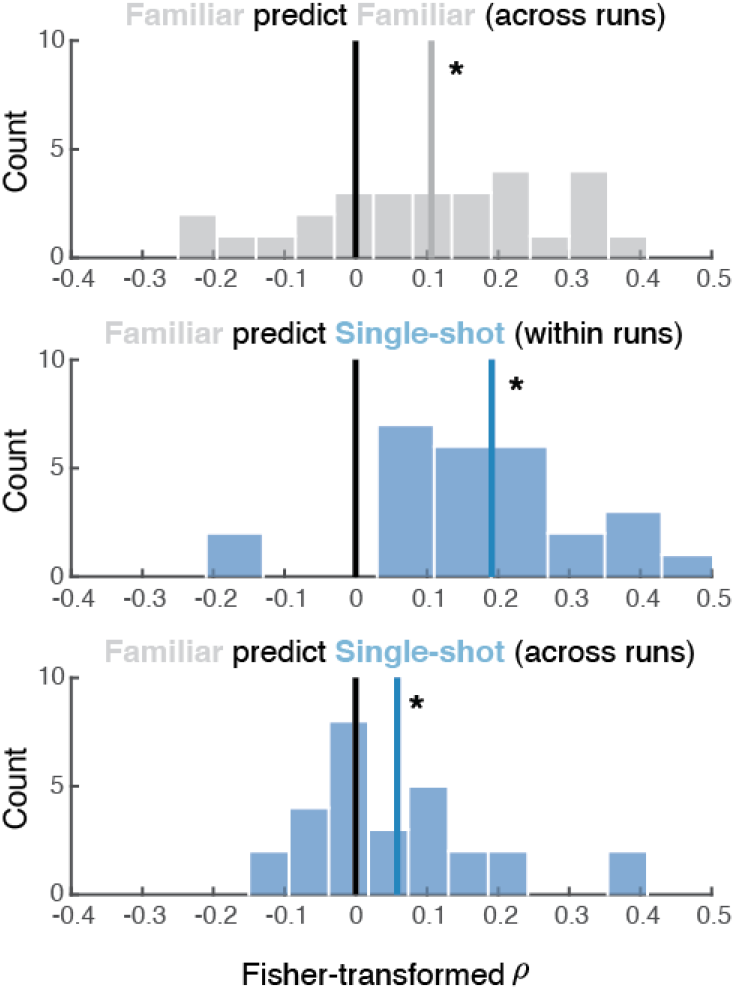
Cross-validated reward prediction error analysis. Regression analyses were run to decode model-defined RPEs from activation in dorsal striatal voxels in the Familiar condition. We used the resulting regression weights at each voxel to generate predicted trial-by-trial RPEs for the held-out runs both within and between conditions. Plots depict the distribution of correlation coefficients between predicted RPEs (derived from BOLD data) with model-derived RPEs. * = *p* < 0.05

**Figure 5.**
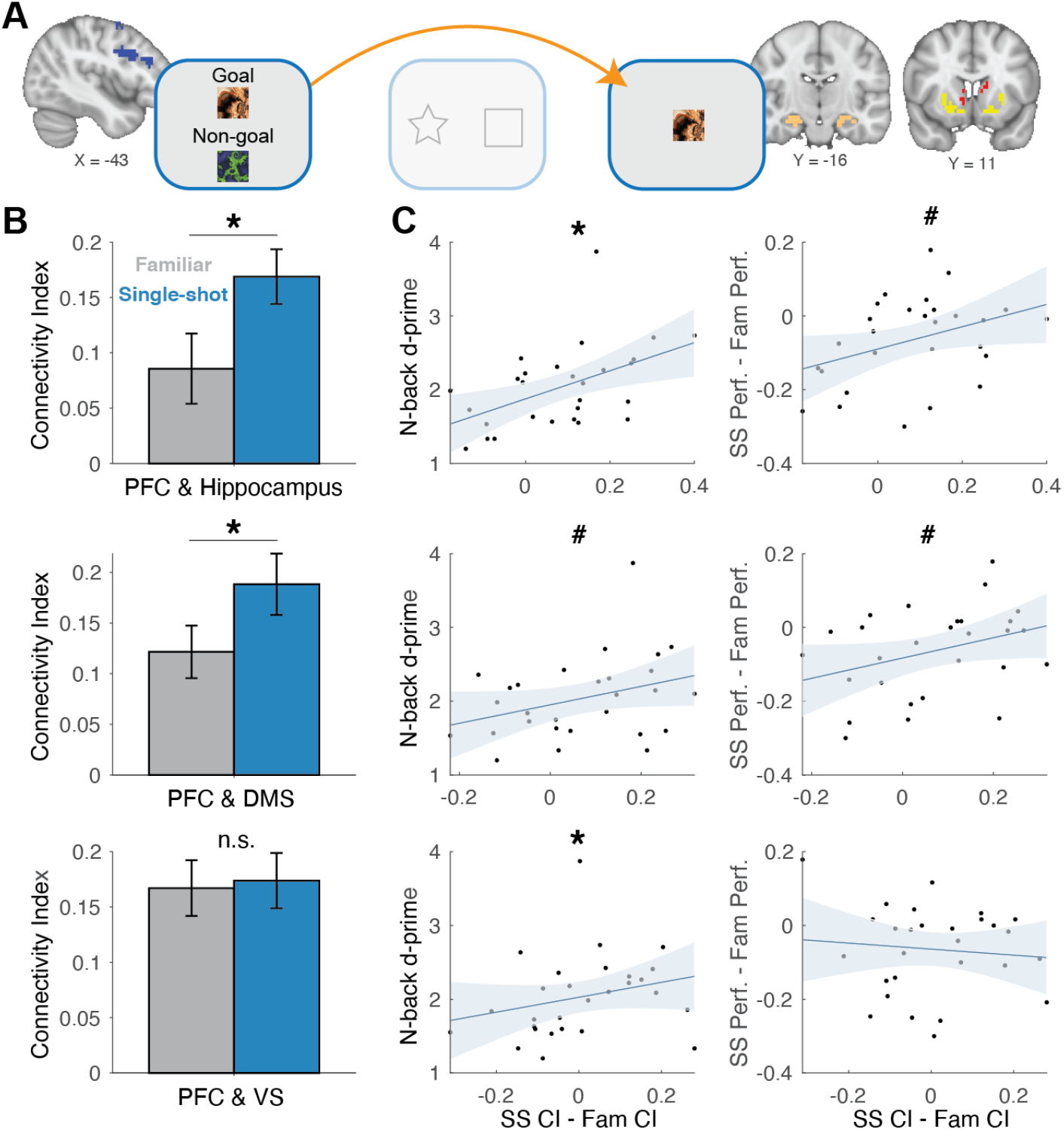
Functional correlations across brain regions and trial phases. **A)** Functional correlations were computed between PFC activity during the pre-choice phase and feedback-locked hippocampal and striatal activity (dorsal and ventral) on rewarded trials. **B)** Within-subject functional correlation results. **C)** Between-subject correlations relating differences in connectivity between conditions, and both n-back performance (left column) and learning task performance (right columns) as a function of condition. CI = connectivity index; SS = Single-shot feedback condition; Fam = Familiar feedback condition; PFC = prefrontal cortex; DMS = dorso-medial striatum; VS = ventral striatum. Shaded regions connote 95% confidence intervals. *#* = *p* < 0.10; * = *p* < 0.05.

As predicted, we found stronger functional correlations in the Single-shot condition between PFC during goal encoding and reward (hippocampus) and RPE-related (DMS) areas at feedback, (PFC-hippocampus, *t*(27) = 3.01, *p* = 0.006; PFC-DMS, *t*(27) = 2.36, *p* = 0.026). We observed no significant condition difference in PFC-VS functional correlations (*t*(27) = 0.25, *p* = 0.81). These results suggest that transient goal encoding in the PFC may drive downstream reward signals in subcortical structures when those goals are attained.

One alternative explanation for this result is that elevated attention or vigilance in the Single-shot condition’s pre-choice phase could drive higher global activity across multiple brain regions that persists into the feedback phase. First, we note that the null PFC-VS result (Figure 5B) speaks against this global confound. Nonetheless, we controlled for this possibility by performing the above connectivity analysis in two additional regions that have been shown to respond to rewards, the posterior cingulate cortex (PCC; McDougle et al., 2019; Pearson et al., 2011) and thalamus (Knutson et al., 2001). We did not expect these ROIs to contribute significantly to the hypothesized goal learning processes. Indeed, there were no significant effects of feedback condition for PFC functional correlations with either the PCC (Figure S5; *t*(27) = 0.71, *p* = 0.48) nor the thalamus (Figure S5; *t*(27) = 0.97, *p* = 0.34), making a global attentional account of our results unlikely.

The observed connectivity results also appeared to be uniquely related to PFC processing during the encoding of goal stimuli (the pre-choice phase), rather than PFC activity during the feedback phase: We observed no significant differences between Single-shot versus Familiar functional correlations between feedback-locked PFC activation and feedback-locked hippocampal, DMS, or VS activation (all *p*s > 0.47). Thus, our results did not appear to be driven by heightened PFC activity that persisted throughout Single-shot trials. Rather, initial encoding of goals (reflected in PFC activation) covaried in a trial-by-trial manner with the strength of the subcortical outcome responses.

We reasoned that covariation between executive and reward regions would be related both to executive functioning itself (n-back performance), and, critically, to endowing novel goals with value to improve learning. Indeed, between subjects, the difference between PFC and hippocampus connectivity strength in the Single-shot versus Familiar condition was significantly correlated with n-back performance (*ρ* = 0.56, *p* = 0.002), and was marginally correlated with Single-shot versus Familiar condition learning differences (*ρ* = 0.37, *p* = 0.052; Figure 5C, top row). Moreover, the degree of difference between PFC and DMS connectivity strength in the Single-shot versus Familiar condition was marginally correlated with both n-back performance (*ρ* = 0.33, *p* = 0.090) and condition learning differences (*ρ* = 0.33, *p* = 0.088; Figure 5C, middle row). We also observed a significant correlation between the difference between PFC and VS connectivity in the Single-shot versus Familiar condition and n-back performance (*ρ* = 0.38, *p* = 0.046; Figure 5C, bottom row), but not corresponding learning differences (*ρ* = 0.10, *p* = 0.62). We note that although the connectivity results were robust in our within-subject contrasts (Figure 5B), the mostly trending between-subjects correlation effects (Figure 5C) should be interpreted with caution. However, taken as a whole, these results suggest that top-down cortical processes may rapidly endow goals with value to shape downstream reward responses and promote adaptive decision-making.

### PFC interactions with dopaminergic regions for learning from single-shot goals

In addition to the planned analyses, we also performed an additional exploratory, *a posteriori* analysis to test if rapidly endowing goals with value relates to interactions between PFC and the ascending dopaminergic system. To test this, we examined functional correlations between our PFC ROI and an anatomically-defined ventral tegmental area (VTA) ROI (Murty et al., 2014), the main source of the brain’s mesolimbic and mesocortical dopamine (Figure 6).

**Figure 6.**
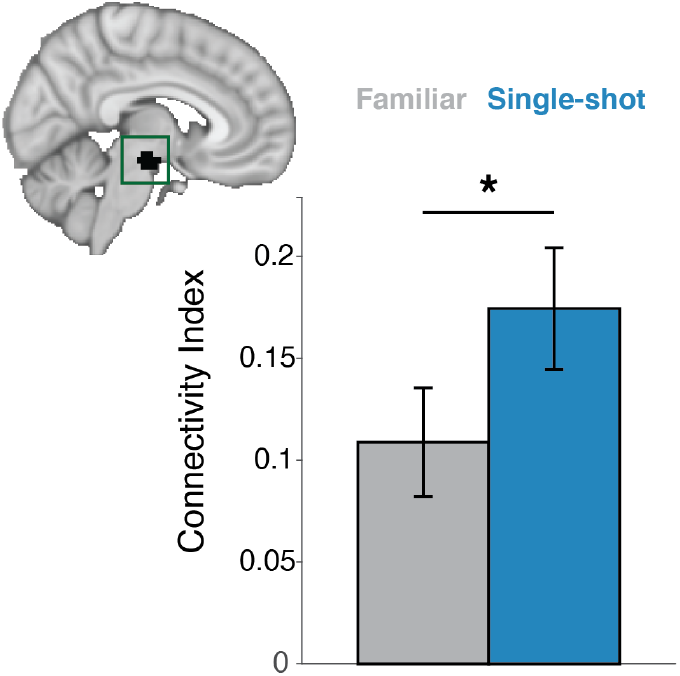
Functional correlations between PFC and the VTA. Functional correlations between PFC activity during the pre-choice phase and feedback-locked ventral tegmental area (VTA) activity on successful trials. Inset: anatomical VTA ROI used in the analysis (shown in black). * = *p* < 0.05.

Functional correlations between PFC (during pre-choice) and VTA (during feedback) were significantly higher in the Single-shot condition relative to the Familiar condition (Figure 6; *t*(27) = 2.31, *p* = 0.029). These effects were unique to PFC activity during the pre-choice phase: Correlations between feedback-locked PFC activity and feedback-locked VTA activity were not significantly different between conditions (*t*(27) = 0.25, *p* = 0.80). However, we did not observe significant between-subjects brain-behavior correlations – PFC-VTA connectivity differences between conditions did not correlate with n-back performance nor learning differences (*p*’s > 0.60). Taken together, these results offer preliminary evidence that single-trial endowment of novel transient goals with value might involve PFC interactions with dopaminergic neurons (Ballard et al., 2011; Sharpe et al., 2019).

## DISCUSSION

Here we presented evidence that learning through attaining transient novel goals is behaviorally similar to learning via familiar secondary reinforcers. This type of learning appears to rely on highly similar activation of reward circuitry. Specifically, we found overlapping neural responses to single-shot goal attainment and familiar reward attainment with respect to outcome valence, suggesting that transient goals can substitute as rewards during instrumental learning. During choice, value representations were similar between conditions, supporting the idea that learning mechanisms were shared. Similarly, reward prediction errors, the key teaching signal of reinforcement learning, appeared to be similar between the single-shot goal and familiar reward conditions, especially in cortical regions. Lastly, successful performance in the single-shot goal condition was associated with increased connectivity between prefrontal cortical regions implicated in executive control and extracortical reward circuits. Together, these findings are consistent with executive function enabling an arbitrarily flexible reward function for corticostriatal reinforcement learning (Daniel & Pollmann, 2014).

The crucial feature of our experimental design was the fact that the goals (i.e., fractal stimuli) were novel on every Single-shot condition trial. Thus, to effectively learn in this condition, subjects needed to rapidly endow value to these transient stimuli (Cole et al., 2013). How does a never-before-seen stimulus get rapidly imbued with value? We propose that a novel stimulus can be internally defined as a goal (e.g., through verbal/symbolic instruction), and then be endowed with value while held in working memory. With these ingredients, attaining this goal would then express the key features of a typical reinforcer. Testing this proposal is difficult, though our results provide some evidence in its favor: Performance on an independent executive function task predicted subjects’ ability to learn from single-shot goals, even when controlling for task difficulty (Figure 2D). Moreover, BOLD activity in prefrontal executive regions during the specification of single-shot goals positively correlated with subsequent responses in the brain’s reward system (Figures 5 and 6).

Important questions remain about the factors that render a novel stimulus valuable within the brain’s reward system. Good performance in the single-shot goal task might require intrinsic motivation (Barto, 2013; Chentanez et al., 2005), where actions are reinforced essentially “for their own sake.” Although the novel goals were extrinsic visual cues, the motivation to learn from them in the first place – even though they lacked any prior value – could simply reflect an intrinsic desire to follow the experimenter’s instructions (Doll et al., 2009), a social learning process that is reinforced over development. Interestingly, a previous study found that dopaminergic medication can boost a learner’s ability to adhere to new instructions about previously learned neutral stimulus–outcome associations that were only endowed with value after learning (Smittenaar et al., 2012). Our findings are consistent with this result, supporting a role for the dopaminergic system in treating abstract goals as rewards via explicit (verbal or symbolic) instructions. Further research could investigate if other independent correlates of intrinsic motivation (Deci, 1971) predict one’s ability to learn from transient novel goals, and engender flexibility in the subcortical reward system.

Our results also add a complication to a strict dichotomy between goal-directed and instrumental components of decision-making (Collins & Cockburn, 2020; Dickinson & Balleine, 1994; (Doll et al., 2012). In our study, we observed essentially overlapping neural signatures for single-shot goal outcomes and secondary reinforcer outcomes, the latter reflecting conventional model-free learning feedback (Figure 3). Other studies have revealed distinct networks corresponding to these two cases: For instance, Gläscher et al. (2010) showed that neural signatures of model-based state prediction errors (in lateral PFC and the intraparietal sulcus) were physiologically distinct from neural signatures of reward prediction errors (in ventral striatum). These signals reflected violations of state transition expectations during a sequence of choices. Gläscher and colleagues’ findings and similar results suggest that subjects’ behavior in our Single-shot condition could in theory be driven by this type of non-RL process. Much like learning a complex trajectory toward a goal, subjects could learn a transition structure between choice stimuli and the “goal fractal” category of objects, without ever needing to rely on a “bottom-up” reward signal. However, our results do not support this particular interpretation, as we observed evidence for clear engagement of typical reward learning circuits during learning from single-shot goals. We propose that the contributions of reward circuits to learning from goals is a function of the task design: If goals are endowed with motivational value, learning should migrate from a state prediction system to a typical reinforcement learning system. Put another way, when the “state” that would normally contribute to a state prediction error reflects goal attainment (as in our task), the line between a state prediction error and a reward prediction may be blurred. Indeed, other studies have revealed similar overlaps between state and reward prediction errors in canonical reinforcement learning circuits (Guo et al., 2016; Langdon et al., 2017).

In contrast to the current study, previous research has highlighted important differences between attaining goals versus attaining rewards during hierarchically organized decision-making. For example, Ribas-Fernandes et al. (2011) used a hierarchical task that included a sub-goal (i.e., pick up a package in a mail delivery simulation) that had to be attained before earning a terminal reward (i.e., successful delivery of the package). The authors observed neural correlates of a pseudo-reward prediction error, driven by surprising “jumps” in the location of the sub-goal, in regions including the anterior cingulate, insula, lingual gyrus, and right nucleus accumbens. However, in a separate behavioral experiment they argued that sub-goal attainment was not a reliable proxy for reinforcement. It is possible that a lack of need for protracted learning from sub-goals in such studies of hierarchical decision-making (Ribas-Fernandes et al., 2011) may lead to qualitatively different neural responses to goals versus studies like ours, where attaining goals is a requirement for learning.

Our study has several important limitations. First, our task design, where novel goals had to be encoded and briefly maintained in short-term memory, may have artificially introduced a dependence on executive function. That is, our task made it somewhat difficult to fully separate effects of learning from novel goals from effects of engaging working memory, as both were clearly required. It should be mentioned that even tasks like the n-back, the striatum shows positive responses on correct detection trials (Satterthwaite et al., 2012), suggesting that performing such tasks correctly can provide an intrinsic reward. Indeed, we replicated this finding in the current data set (Figure S6). There are, however, cases where a novel goal could be acted on without requiring working memory: For instance, a goal could instead be retrieved from episodic memory, such as recalling and acting on an instruction you received hours (or days) in the past. In that case, the medial temporal lobe and medial prefrontal cortex may be involved in maintaining and communicating the features of valuable goals to striatal and midbrain circuits (Han et al., 2010). Either way, it is difficult to conceive of a setting where learning from novel goals would not carry memory demands, whether short-or long-term.

Second, the precise cause of the poorer performance we observed in the Single-shot feedback condition (Figure 2A) was not clear. Our modeling analysis appeared to rule out the interpretation that the effect was driven by noise during choice. Although it was apparent that executive function – operationalized by performance in the independent n-back task – was selectively related to learning in the Single-shot goal condition (Figure 2D), multiple psychological phenomena could have caused attenuated performance in that condition. These include a weaker appetitive signal for single-shot goal outcomes, or an increased frequency of lapses of attention, where the goal fractal is either not encoded initially or forgotten by the time of feedback. Future behavioral studies could attempt to fill this gap, for example by testing subjects’ memory of the goal images themselves on intermittent probe trials or in a subsequent long-term memory test.

Third, we found mixed results with respect to striatal reward prediction error (RPE) encoding in the Single-shot condition (Figures 3 and 4). Surprisingly, our cross-validated encoding analysis (Figure 4) supported the presence of striatal RPEs in the Single-shot condition, while our more liberal beta calculation showed non-significant RPE encoding in the Single-shot condition (Figure 3B, Figure S3). These inconsistent results could suggest a lack of power or sensitivity, or a true attenuation of prediction errors when they are signaled by goals held in working memory. The latter interpretation would be consistent with findings showing a weakening of striatal prediction error signals when working memory is contributing to choice behavior (Collins et al., 2017). One approach to address this question could be to induce a wider dynamic range of RPEs, or match working memory demands.

Learning from goals in addition to familiar reinforcers is a key aspect of intelligent behavior, and is an especially important cognitive tool for humans. Here, we pushed this concept to its logical extreme, asking if goals can stand in for rewards during learning even when those goals are abstract stimuli with no prior meaning or value to the learner, and are only observed a single time. We demonstrated that human subjects can easily perform this kind of single-shot instrumental learning, and that this form of learning shares many behavioral and physiological features with conventional instrumental learning from secondary reinforcers. The ability to rapidly direct behavior towards the completion of goals has been linked to executive control processes in the human prefrontal cortex (Duncan et al., 1996), a finding that our data further supports. Taken together, our findings suggest that humans can rapidly and flexibly define what constitutes a reinforcer in a single instance, harnessing the brain’s executive functions and reward circuitry to optimize goal-directed behavior.

## METHODS

### Participants

Thirty-two healthy volunteers (aged 18-40; mean age = 25.6 years; 18 females) participated in the experiment. All subjects were right-handed, had no known neurological disorders, and had normal or corrected-to-normal vision. Subjects provided written, informed consent, and received $30 for their participation in the experiment. Functional neuroimaging data from three subjects were not analyzed beyond preprocessing because of excessive head motion (see below for details on these exclusion criteria), and an additional subject was excluded from both behavioral and neural analyses for showing overall below-chance mean performance in the task (i.e., more often choosing the less-rewarding stimulus). The three subjects excluded for excessive head motion were included in the behavioral and modeling analyses, yielding an effective sample size of 31 subjects for behavior analysis; the effective sample size for imaging analysis was 28 subjects. Experimental protocols were approved by the Institutional Review Board at the University of California, Berkeley.

### Probabilistic Selection Task

Subjects were tasked with learning which of two stimuli, across four pairs of stimuli, was more likely to yield a favorable outcome (Figure 1A). For one run, the choice stimuli were black line-drawings of simple shapes (e.g., diamond, circle, triangle, etc.), and for the other run they were differently-colored squares (e.g., blue, red, green, etc.). The order of these two runs was counterbalanced across subjects. Stimuli were presented using MATLAB software (MathWorks) and the Psychophysics Toolbox (Brainard, 1997). The display was projected onto a translucent screen that subjects viewed via a mirror mounted on the head coil.

Trials proceeded as follows (Figure 1A). First, during the pre-choice phase, the type of feedback associated with the current trial was displayed (see below for details). This phase lasted 2 s. After a brief inter-stimulus interval (0.5-2.5 s, uniform jitter), the choice phase began, and a single pair of choice stimuli were presented (e.g., square versus circle). The sides (left or right of the central fixation cross) where each of the two stimuli appeared was randomized across trials. Subjects had 1.2 s to render their choice with an MR-compatible button box, using their index finger to select the stimulus on the left and their middle finger to select the stimulus on the right. Successfully registered choices were signaled by the central fixation cross changing from black to blue. The choice phase ended after 1.2-2 s (uniform jitter) regardless of the reaction time, and choice stimuli stayed on screen until a second inter-stimulus interval with only the fixation cross displayed (0.5 s). Finally, in the feedback phase, feedback was presented for 1 s, followed by an inter-trial interval of 1.5-3 s (uniform jitter). If reaction times were too fast (< 0.2 s) or too slow (> 1.2 s), the trial was aborted and a “Too Fast!” or “Too Slow!” message (red font) was displayed centrally for 1 s in lieu of any reward feedback and the ITI was initiated. (4.12 ± 0.73% of trials were aborted in this manner; mean ± 95% s.e.m.).

Two reward conditions were used (Familiar versus Single-shot; Figure 1A) as well as two difficulty levels (Easy versus Hard; Figure 1B). In the Familiar condition, feedback “point” stimuli were previewed during the pre-choice phase, with the “+1” presented toward the top of the display and the “+0” presented toward the bottom, accompanied by the text “POINTS trial” in black font. At feedback, rewarded choices were signaled by numeric points, where a “+1” message was presented on successful trials (green font), and a “+0” message was presented on unsuccessful trials (red font). In the Single-shot condition pre-choice phase, two distinct colored fractal-like patterns were displayed, with one labeled as the “Goal” (displayed toward the top of the screen) and the other as the “Non-goal” (displayed toward the bottom of the screen), accompanied by the text “GOAL trial” in black font. Here, the subject’s task was to remember the identity of the goal fractal, so that in the feedback phase they can appropriately reinforce their preceding choice of stimulus. Colorful fractal stimuli were created using the randomize function of ArtMatic Pro (www.artmatic.com), and the final set of fractals was selected to maximize discriminability between any two fractals.

Trials were also evenly divided into two levels of difficulty: Easy or Hard. On Easy trials, the optimal choice stimulus yielded a favorable outcome (i.e., +1 point or the goal fractal) on 90% of trials (27 of 30 trials), and an unfavorable outcome (i.e., +0 points or the non-goal fractal) on 10% of trials. The other choice stimulus on Easy trials was associated with the inverse success probability (10%/90%). On Hard trials, the optimal choice stimulus yielded a favorable outcome (i.e., one point or the goal fractal) on ∼77% of trials (23 of 30 trials), and an unfavorable outcome (i.e., 0 points or the non-goal fractal) on ∼23% of trials. The other choice stimulus on Hard trials was associated with the inverse success probability (23%/77%). Reward schedules were deterministically pre-specified so that the exact number of successful/unsuccessful outcomes for each stimulus was identical across subjects, though each subject received a unique pseudo-randomized trial sequence. Crossing the two conditions (Familiar and Single-shot) and two difficulty levels (Easy and Hard) yielded four choice stimulus pairs for each run. Subjects performed 120 trials in each of the two runs of the task, performing 30 choices per run for each condition (i.e., for each of the four pairs of choice stimuli). To ensure that subjects understood the task, the experiment began with a thorough explanation of the conditions followed by a sequence of 16 practice trials that included both Familiar and Single-shot trial types (with deterministic reward/goal feedback), using goal fractal images and choice stimuli (Klingon characters) that were not seen in either subsequent learning run. These practice trials occurred during the anatomical scan.

In each run, after learning trials were completed, a “surprise” testing phase was administered (we note that the surprise aspect only existed for the first run of the experiment). Here, after a brief break cued by an instruction screen (12 s), a pseudo-random sequence of choice stimulus pairs was presented (1 s each, max RT 1 s, inter-trial interval 1.5-3 s, uniform jitter). In this phase all possible pairings of the 8 choice stimuli seen during learning were presented three times each, yielding 84 total trials. Participants made choices in the testing phase, though no feedback was given.

### N-back task

After completing both learning runs, subjects also performed an n-back task during the third and final functional scan. In this task, subjects were shown a pseudo-randomized sequence of novel opaque black shapes (ten unique stimuli; stimuli source: Vanderplas & Garvin, 1959; task code modified from https://github.com/JAQuent/nBack), and asked to respond with a button press (forefinger press on the MR-compatible button box) whenever a shape repeated following *N* intervening different shapes. The current *N-*rule was specified via an instruction screen at the start of each sequence.

Four sequences (blocks) were performed at each *N* (Ns used: 1, 2, 3). Each sequence was 17 + *N* items long, had 7 target trials (hits) per sequence, and had no triple repeats of any single shape nor any repeats in the first three presented shapes. Each shape appeared centrally for 0.5 s, with a 1.5 s inter-stimulus interval. A black fixation cross was presented throughout the sequences. The order of the twelve sequences was randomized, although the first three sequences of the task were fixed to *N* = 1, 2, 3 in that order.

### Behavioral analysis

Behavior during the learning phase of the probabilistic selection task was quantified by a simple performance metric that reflected the percent of trials where subjects chose the optimal stimulus in each pair (i.e., the stimulus most likely to yield a reward; Figure 1; Frank et al., 2007). In the testing phase, behavior was quantified using a logistic regression model. In this model, the response variable was a boolean based on whether the subject chose the stimulus on the right side of the screen (1) or the left side (0). Predictors included the cumulative reward difference of the stimuli (right minus left) determined by the sum of rewards (points or fractals) yielded by that stimulus during the preceding learning phase; a boolean predictor for trials that pitted two Familiar condition stimuli against one another; a boolean predictor for trials that pitted two Single-shot condition stimuli against one another; and a signed predictor capturing a Familiar condition bias, with a value of 1 when a Familiar condition stimulus appeared on the right and an Single-shot condition stimulus appeared on the left, a value of −1 for when an Single-shot condition stimulus appeared on the right and a Familiar condition stimulus appeared on the left, and a value of 0 when both stimuli were associated with the same condition. Interaction terms were included as well, and all predictors were z-scored.

N-back performance was quantified using the d-prime metric (Haatveit et al., 2010). Correlations between n-back performance (d-prime over all trials/Ns) and performance in the probabilistic selection task were computed using both Spearman and Pearson correlations, and were conducted on both the full set of trials in the probabilistic selection task, or the subset that showed similar cross-condition performance (i.e., Familiar-Hard trials and Single-shot-Easy trials; Figure 1D).

### Computational modeling analysis

We tested several variants of standard trial-based reinforcement learning (RL) models (Sutton & Barto, 1998) to account for subjects’ instrumental learning behavior and to build model-derived regressors for fMRI analyses. All models tested were built using the same basic architecture, where values of stimuli were updated according to the delta rule:

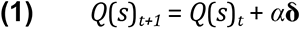

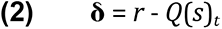

Here *Q*(*s*)*_t_* reflects the learned value of stimulus *s* on trial *t*, *α* reflects the learning rate, and **δ** reflects the reward prediction error (RPE), or the difference between the observed reward (*r*) and the expected reward given the choice of stimulus *s*. Action selection between the two presented stimuli was modeled using the softmax function,

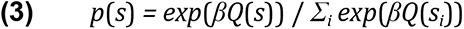

Where *β* is the inverse temperature parameter.

Two candidate models also included a decay parameter, which captured trial-by-trial forgetting of all stimulus *Q*-values,

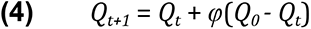

Where 0 < *φ* < 1 is a decay parameter which at each trial pulls value estimates towards their initial value *Q_0_* = 0. (We note that a preliminary analysis found that the best-fitting value of initial *Q*-values, *Q_0_*, was 0).

After performing a preliminary model fitting and comparison analysis across a wide range of candidate models (using maximum likelihood estimation), we narrowed our primary model fitting and comparison analysis to six candidate model variants that performed well in our first-pass analysis and represented distinct behavioral interpretations (see Table S1 for details of each tested model).

The first model we tested reflected our hypothesis that the differences in performance we observed between the Familiar and Single-shot feedback conditions were strictly a function of weaker learning in the Single-shot condition. We implemented this by having an independent learning rate (*α*) for each feedback condition (Familiar versus Single-shot). This model included a single decay parameter and a single temperature parameter.

In a second variant we assumed that performance differences were a function of both weaker learning and increased forgetting of stimulus values in the Single-shot condition; this model matched the first model but had unique decay parameters for each feedback condition. In a third variant we excluded the decay parameter altogether but included feedback condition-specific learning rates, and in a fourth variant we assumed asymmetric updates (i.e., unique learning rates) for positive and negative outcomes (Frank et al., 2007; Gershman, 2015)

To capture a form of rapid learning that is qualitatively distinct from incremental RL, we also tested a simple heuristic “win-stay lose-shift” model, which can be captured by Equation 1 when *α* = 1. To allow for feedback condition differences in this simple heuristic model, we included a unique *β* free parameter for each feedback condition.

Finally, in our sixth model we tried to capture the hypothesis that the learning processes (Equations 1 and 2) in both the Familiar and Single-shot condition were identical (via equal learning rates), but that the choice process (Equation 3) was different, perhaps being noisier in the Single-shot condition due to the presence of a secondary task (i.e., having to maintain the goal fractal during the choice phase). We approximated the effect of putative choice-phase noise via an undirected noise parameter,

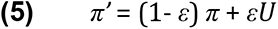

Here, given the action selection policy *π* = *p*(*s*) (Equation 3), Equation 5 introduces noise by implementing a mixed policy where *U* is the uniform random policy (i.e., a coin flip between stimuli) and the parameter 0 < *ε* < 1 controls the amount of decision noise introduced during choice (Collins et al., 2014). We included separate *ε* free parameters for each feedback condition.

For model fitting and model comparisons, we used a recently-developed technique, Hierarchical Bayesian Inference (Piray et al., 2019). In this method (HBI), models are compared and free parameters are estimated simultaneously. Recent work using this method has shown that fits estimated by HBI are more precise and recoverable than competing methods, and model comparison is robust and less biased towards simplicity (Piray et al., 2019). We note briefly that similar behavioral and fMRI results were obtained when we used traditional maximum likelihood estimation methods; however, clearer model comparison results and more interpretable parameter estimates were obtained in our HBI analysis. For our HBI procedure we implemented a prior variance parameter of *v* = 6.00. Finally, to compare learning rate parameters across feedback conditions (Figure 1F) we could not perform standard frequentist tests due to the hierarchical procedure; thus, in a follow-up HBI analysis we implemented the Single-shot condition learning rate as a free parameter that was added or subtracted to the Familiar condition learning rate and performed an “HBI t-test” on the the resulting parameter fit relative to zero. (Further details concerning the HBI analysis procedure, and its open-source toolbox, are given in Piray et al., 2019.)

After model fitting and comparison, we validated the winning model by simulating choice behavior using the best-fit parameters. For each subject, we simulated 100 RL agents using the schedule of rewards seen by that subject and the best-fit individual-level parameters for that subject gleaned from the HBI procedure. Results were averaged before plotting (Figure 2A, B). For model-derived fMRI analyses, we simulated the model for each subject using their actual choice history and best-fit learning rate and decay parameters. This procedure yielded model-based predictions of trial-by-trial reward prediction errors (RPEs) and trial-by-trial *Q-*values (of the chosen stimulus). Both of these variables were convolved with the canonical hemodynamic response function and used to model BOLD responses during, respectively, the feedback and choice phases. Lastly, in a control fMRI analysis, we instead used the group-level parameters for each subject (as given by the HBI fitting procedure) to simulate RPEs (Figure S3).

### Imaging procedures

Whole-brain imaging was performed at the Henry H. Wheeler Jr. Brain Imaging Center at the University of California, Berkeley on a Siemens 3T *Trio* MRI scanner using a 12-channel head coil. Functional data were acquired with a gradient-echo echo-planar pulse sequence (TR = 2.25 s, TE = 33 ms, flip angle = 74°, 30 slices, voxel size = 2.4 mm x 2.4 mm x 3.0 mm). T1-weighted MP-RAGE anatomical images were collected as well (TR = 2.30 s, TE = 2.98 ms, flip angle = 9°, 160 slices, voxel size = 1.0 mm isotropic). Functional imaging was performed in three runs, with the first two runs consisting of the probabilistic selection task (584 volumes each) and the third run consisting of the n-back task (275 volumes). A field map scan was performed between the two probabilistic selection task runs to correct for magnetic field inhomogeneities (see *Image Preprocessing*). Subjects’ head movements were restricted using foam padding.

### Image preprocessing

Preprocessing was performed using fMRIPrep 1.4.0 (Esteban et al., 2019). First, the T1-weighted (T1w) image was corrected for intensity non-uniformity with N4BiasFieldCorrection (Tustison et al., 2010) and then used as the T1w reference image throughout the workflow. The T1w reference image was skull-stripped (using antsBrainExtraction.sh), and brain tissue segmentation was performed on the brain-extracted T1w using the FSL tool FAST. Brain surfaces were reconstructed using the FreeSurfer tool Recon-all (FreeSurfer 6.0.1; Dale et al., 1999). Volume-based spatial normalization to standard (MNI) space was performed through nonlinear registration with ANTs, using brain-extracted versions of both the T1w reference image and the T1w template.

The functional data were resampled into standard space (MNI), generating a preprocessed BOLD time series for each run. A reference volume (average) and its skull-stripped version (using ANTs for stripping) were generated. A B0 field map was co-registered to the BOLD reference image. Head-motion parameters were estimated with respect to the BOLD reference image (transformation matrices and six corresponding rotation and translation parameters) before spatiotemporal filtering was applied with MCFLIRT (FSL 5.0.9; Jenkinson et al., 2002). Slice-time correction was applied using 3dTshift from AFNI (Cox & Hyde, 1997). The BOLD time-series (including slice-timing correction) were resampled into their original, native space by applying a single transform to correct for head-motion and susceptibility distortions (Glasser et al., 2013). The unwarped BOLD reference volume for each run was co-registered to the T1w reference image using bbregister (FreeSurfer), with nine degrees of freedom to account for any distortions remaining in the BOLD reference image.

Framewise displacement was calculated for each functional run (following Power et al., 2014). Confound regressors for component-based noise correction were created using CompCor (Behzadi et al., 2007). Gridded (volumetric) resamplings were performed using antsApplyTransforms (ANTs), with Lanczos interpolation. Lastly, data were high-pass filtered (100s) and spatially smoothed with a Gaussian kernel (4.0 mm FWHM) prior to all GLM analyses.

### Imaging analyses

Analyses involved five separate GLMs fit to learning phase BOLD data, and one GLM fit to the n-back task BOLD data. All GLM analyses were performed using FSL (version 6.0.3). Regressors of no interest were entered into the model; these included subject reaction time (a parametric regressor yoked to choice onset, convolved with the HRF), button press events (convolved with the HRF), six standard motion regressors, the framewise displacement time course, linear drift terms, and the first 6 aCompCor components. In addition, we added stick regressors corresponding to volumes with large motion artifacts (>1.0 mm framewise displacement, or >5 scaled variance as determined by SPM’s *tsdiffana* protocol), with additional confound stick regressors for the neighboring (subsequent) volume. This procedure was also used to determine movement-related subject exclusions: If more than 10% of volumes in either of the two learning runs were flagged as outliers, the subject was excluded (3 subjects were excluded based on these criteria).

In the first GLM (GLM 1), regressors of interest included condition-specific spike regressors for the onset of each of the following trial phases: pre-choice, choice, feedback, and the choice phase in the testing trials. A feedback-locked valence regressor also was included, which effectively coded successful (+1) and unsuccessful (−1) trials within the learning phase. A model-derived parametric regressor was also included at feedback onset, which captured condition-specific reward prediction errors (RPEs; Equation 2). Including both valence and RPE regressors is important because otherwise brain responses distinguishing the feedback valence (in visual, affective, or cognitive properties) could be misidentified as parametric reward prediction errors. Each subject’s RPE time course was determined using their individually-fit parameters. Orthogonalization was not used, and each regressor was convolved with the canonical hemodynamic response function (double-gamma). Secondary GLMs were also run for two control analyses (Figure S3): In one variant of GLM 1, the RPE regressor in the model was created using group-level learning rate and decay parameter values rather than individually-fit parameters for each subject’s RPE time-course; in a second variant of GLM 1 we included separate positive and negative RPE regressors for each condition. The second GLM (GLM 2) was identical to the first, except that instead of including RPE regressors it included the model-derived *Q*-value for the chosen stimulus as a parametric modulation of choice onset (modeled separately for the two feedback conditions).

The third and fourth GLMs were designed to facilitate decoding and connectivity analyses: GLM 3 included identical task and confound regressors as those in GLM 1, however, feedback onset was modeled on a trial-by-trial basis, with unique stick regressors at each instance of feedback (and no regressors for valence/RPEs). This method produced individual feedback phase beta-maps for each trial of the learning task (Mumford et al., 2014). GLM 4 was also similar to GLM 1, except a single feedback onset regressor for all trials was used, and we instead modeled the pre-choice phase on a trial-by-trial basis, producing individual pre-choice phase beta-maps for each trial of the task.

The n-back task GLM (GLM 5) included the same confound regressors as the learning task GLMs. Regressors of interest included a block-level regressor spanning the beginning to end of each stimulus sequence, and a parametric modulation of that regressor that reflected the particular *N* assigned to each sequence. This latter regressor thus captured the linear effect of increasing cognitive load in the n-back task, and was used as an executive function ROI localizer for later analyses.

Individual subject runs were combined in a fixed effects analysis and then brought to the group level for mixed effect analyses. For whole-brain analyses we imposed a family-wise error cluster-corrected threshold of p < 0.05 (FSL FLAME 1) with a cluster-forming threshold of p < 0.001.

Our region-of-interest (ROI) creation procedure was designed to both avoid double-dipping at the subject level (Boorman et al., 2013; Kriegeskorte et al., 2009) and to conservatively test our predictions about RL processes specifically in the Single-shot feedback condition. First, for each subject we performed a leave-one-out group-level mixed effects analysis of the main effect of interest (e.g., valence) in the Familiar condition trials only, excluding that subject’s data. Pre-threshold masking was performed in this analysis to constrain results within an anatomical region using a cluster-corrected threshold of p < 0.05 (FSL FLAME 1) and cluster-forming threshold p < 0.01 (note that this threshold was relaxed relative to the whole-brain threshold for ROI creation purposes only). For striatal analyses, we used a three-partition striatal mask for anatomical masking (Choi et al., 2012). For cortical ROI masking, we used the Harvard-Oxford probabilistic atlas, thresholding all masks at 0.50. (We note that for the inferior parietal lobule mask we combined the bilateral angular and supramarginal gyri.) Because of this method, resulting Familiar and Single-shot condition ROI results were statistically valid at the subject level, and all Single-shot condition results were additionally validated out-of-set at the condition level. All neural effect sizes (betas) were extracted using *featquery* (FSL).

Our cross-validated encoding analysis of RPE processing (Figure 4) was performed as follows: beta series data for feedback onset (GLM 3) were extracted from each subject’s dorso-medial striatum ROI. Then, for each individual run, a linear regression was performed independently for each voxel, relating its activation to model-derived RPEs on rewarded Familiar condition trials only. We constrained this analysis to rewarded trials to model parametric prediction error effects independent of valence. The resulting *n* X 1 vector of parameter estimates, for *n* voxels, was then multiplied by one of three different *trial* X *n* matrices of independent beta series data: the beta series of rewarded Familiar trials in the held-out run, the beta series of rewarded Single-shot trials within-run, and the beta series of rewarded Single-shot trials in the held-out run. This produced a vector of “predicted” RPEs for each of these analyses. Predicted RPEs were then correlated (Spearman) with the model-derived RPEs for the associated run/condition and Fisher-transformed for statistical analysis (two-tailed *t*-tests relative to zero, with alpha set to 0.05).

Functional correlation analyses were performed as follows: First, we aimed to identify prefrontal cortex (PFC) voxels that were related to the encoding of novel goal stimuli. Thus, we created a PFC ROI using a group-level mixed-effects analysis on the Single-shot>Familiar pre-choice phase contrast (using the aforementioned leave-one-out procedure to maintain statistical independence), masking those results (pre-threshold) with the group main effect map from the n-back GLM (parametric *N* regressor; see above). Then, for each ROI (e.g., PFC, hippocampus, etc.) and phase (e.g., pre-choice, feedback), we extracted trial-by-trial betas from either GLM 3 (for feedback) or GLM 4 (for pre-choice) and averaged within each ROI. We performed beta series correlations (Rissman et al., 2004) by computing the spearman correlation coefficients between each ROI pair, and then Fisher-transformed those values to derive a “Connectivity Index.” We measured both within-subject effects in this Connectivity Index (i.e., Single-shot versus Familiar), as well as between-subjects correlations between the condition differences in connectivity and the condition differences in learning, and n-back performance. Connectivity analyses in control ROIs (thalamus and posterior cingulate cortex; Figure S5) were performed using beta-series data extracted from anatomically-derived ROIs (Harvard-Oxford atlas; thresholded at 0.50). Finally, a *post-hoc* connectivity analysis on PFC-VTA functional correlations (Figure 6) was performed using the probabilistic VTA atlas given by Murty et al. 2014 (thresholded at 0.50).

## ACKNOWLEDGEMENTS

We would like to thank the CCN Lab (UC Berkeley) and ACT Lab (Yale) for helpful discussions. SDM was funded by NIH fellowship F32 MH119797, AGEC and BB are funded by NIH grant R01MH119383, SB is funded by NIH grant R01MH124108, IB is funded by NIH fellowship F32MH119796. This research was supported by a Hellman Fellows Fund award.

## REFERENCES

Babayan, B. M., Uchida, N., & Gershman, S. J. (2018). Belief state representation in the dopamine system. Nature Communications, 9(1), 1891. https://doi.org/10.1038/s41467-018-04397-0

Ballard, I. C., Murty, V. P., Carter, R. M., MacInnes, J. J., Huettel, S. A., & Adcock, R. A. (2011). Dorsolateral Prefrontal Cortex Drives Mesolimbic Dopaminergic Regions to Initiate Motivated Behavior. Journal of Neuroscience, 31(28), 10340–10346. https://doi.org/10.1523/JNEUROSCI.0895-11.2011

Barron, H. C., Dolan, R. J., & Behrens, T. E. J. (2013). Online evaluation of novel choices by simultaneous representation of multiple memories. Nature Neuroscience, 16(10), 1492–1498. https://doi.org/10.1038/nn.3515

Barto, A. G. (2013). Intrinsic Motivation and Reinforcement Learning. In G. Baldassarre & M. Mirolli (Eds.), Intrinsically Motivated Learning in Natural and Artificial Systems (pp. 17–47). Springer. https://doi.org/10.1007/978-3-642-32375-1_2

Bartra, O., McGuire, J. T., & Kable, J. W. (2013). The valuation system: A coordinate-based meta-analysis of BOLD fMRI experiments examining neural correlates of subjective value. NeuroImage, 76, 412–427. https://doi.org/10.1016/j.neuroimage.2013.02.063

Behzadi, Y., Restom, K., Liau, J., & Liu, T. T. (2007). A component based noise correction method (CompCor) for BOLD and perfusion based fMRI. NeuroImage, 37(1), 90–101. https://doi.org/10.1016/j.neuroimage.2007.04.042

Boorman, E. D., Rushworth, M. F., & Behrens, T. E. (2013). Ventromedial Prefrontal and Anterior Cingulate Cortex Adopt Choice and Default Reference Frames during Sequential Multi-Alternative Choice. Journal of Neuroscience, 33(6), 2242–2253. https://doi.org/10.1523/JNEUROSCI.3022-12.2013

Brainard, D. H. (1997). The psychophysics toolbox. Spatial Vision, 10, 433–436.

Charpentier, C. J., Bromberg-Martin, E. S., & Sharot, T. (2018). Valuation of knowledge and ignorance in mesolimbic reward circuitry. Proceedings of the National Academy of Sciences, 115(31), E7255–E7264. https://doi.org/10.1073/pnas.1800547115

Chentanez, N., Barto, A. G., & Singh, S. P. (2005). Intrinsically Motivated Reinforcement Learning. In L. K. Saul, Y. Weiss, & L. Bottou (Eds.), Advances in Neural Information Processing Systems 17 (pp. 1281–1288). MIT Press. http://papers.nips.cc/paper/2552-intrinsically-motivated-reinforcement-learning.pdf

Choi, E. Y., Yeo, B. T. T., & Buckner, R. L. (2012). The organization of the human striatum estimated by intrinsic functional connectivity. Journal of Neurophysiology, 108(8), 2242–2263. https://doi.org/10.1152/jn.00270.2012

Cole, M. W., Laurent, P., & Stocco, A. (2013). Rapid instructed task learning: A new window into the human brain’s unique capacity for flexible cognitive control. Cognitive, Affective, & Behavioral Neuroscience, 13(1), 1–22. https://doi.org/10.3758/s13415-012-0125-7

Collins, A. G., Brown, J. K., Gold, J. M., Waltz, J. A., & Frank, M. J. (2014). Working memory contributions to reinforcement learning impairments in schizophrenia. Journal of Neuroscience, 34(41), 13747–13756.

Collins, A. G. E. (2018). The Tortoise and the Hare: Interactions between Reinforcement Learning and Working Memory. Journal of Cognitive Neuroscience, 30(10), 1422–1432. https://doi.org/10.1162/jocn_a_01238

Collins, A. G. E., Ciullo, B., Frank, M. J., & Badre, D. (2017). Working Memory Load Strengthens Reward Prediction Errors. The Journal of Neuroscience, 37(16), 4332–4342. https://doi.org/10.1523/JNEUROSCI.2700-16.2017

Collins, A. G. E., & Cockburn, J. (2020). Beyond dichotomies in reinforcement learning. Nature Reviews Neuroscience, 21(10), 576–586. https://doi.org/10.1038/s41583-020-0355-6

Collins, A. G. E., & Frank, M. J. (2013). Cognitive control over learning: Creating, clustering, and generalizing task-set structure. Psychological Review, 120(1), 190–229. https://doi.org/10.1037/a0030852

Collins, A. G. E., & Frank, M. J. (2018). Within- and across-trial dynamics of human EEG reveal cooperative interplay between reinforcement learning and working memory. Proceedings of the National Academy of Sciences, 115(10), 2502–2507. https://doi.org/10.1073/pnas.1720963115

Cowles, J. T. (1937). Food-tokens as incentives for learning by chimpanzees (p. 96). The Johns Hopkins Press. https://doi.org/10.1037/14268-000

Cox, R. W., & Hyde, J. S. (1997). Software tools for analysis and visualization of fMRI data. NMR in Biomedicine, 10(4–5), 171–178. https://doi.org/10.1002/(SICI)1099-1492(199706/08)10:4/5<171::AID-NBM453>3.0.CO;2-L

Dale, A. M., Fischl, B., & Sereno, M. I. (1999). Cortical Surface-Based Analysis: I. Segmentation and Surface Reconstruction. NeuroImage, 9(2), 179–194. https://doi.org/10.1006/nimg.1998.0395

Daniel, R., & Pollmann, S. (2010). Comparing the Neural Basis of Monetary Reward and Cognitive Feedback during Information-Integration Category Learning. Journal of Neuroscience, 30(1), 47–55. https://doi.org/10.1523/JNEUROSCI.2205-09.2010

Daniel, Reka, & Pollmann, S. (2014). A universal role of the ventral striatum in reward-based learning: Evidence from human studies. Neurobiology of Learning and Memory, 114, 90–100. https://doi.org/10.1016/j.nlm.2014.05.002

Davidow, J. Y., Foerde, K., Galván, A., & Shohamy, D. (2016). An Upside to Reward Sensitivity: The Hippocampus Supports Enhanced Reinforcement Learning in Adolescence. Neuron, 92(1), 93–99. https://doi.org/10.1016/j.neuron.2016.08.031

Daw, N. D., Gershman, S. J., Ben Seymour, Dayan, P., & Dolan, R. J. (2011). Model-Based Influences on Humans’ Choices and Striatal Prediction Errors. Neuron, 69(6), 1204–1215. https://doi.org/10.1016/j.neuron.2011.02.027

Daw, N. D., O’Doherty, J. P., Dayan, P., Seymour, B., & Dolan, R. J. (2006). Cortical substrates for exploratory decisions in humans. Nature, 441(7095), 876–879. https://doi.org/10.1038/nature04766

Deci, E. L. (1971). Effects of externally mediated rewards on intrinsic motivation. Journal of Personality and Social Psychology, 18(1), 105–115. https://doi.org/10.1037/h0030644

Delgado, M. R., Nystrom, L. E., Fissell, C., Noll, D. C., & Fiez, J. A. (2000). Tracking the Hemodynamic Responses to Reward and Punishment in the Striatum. Journal of Neurophysiology, 84(6), 3072–3077. https://doi.org/10.1152/jn.2000.84.6.3072

Dickinson, A., & Balleine, B. (1994). Motivational control of goal-directed action. Animal Learning & Behavior, 22(1), 1–18. https://doi.org/10.3758/BF03199951

Doll, B. B., Jacobs, W. J., Sanfey, A. G., & Frank, M. J. (2009). Instructional control of reinforcement learning: A behavioral and neurocomputational investigation. Brain Research, 1299, 74–94. https://doi.org/10.1016/j.brainres.2009.07.007

Doll, B. B., Simon, D. A., & Daw, N. D. (2012). The ubiquity of model-based reinforcement learning. Current Opinion in Neurobiology, 22(6), 1075–1081. https://doi.org/10.1016/j.conb.2012.08.003

Duncan, J., Burgess, P., & Emslie, H. (1995). Fluid intelligence after frontal lobe lesions. Neuropsychologia, 33(3), 261–268. https://doi.org/10.1016/0028-3932(94)00124-8

Duncan, J., Emslie, H., Williams, P., Johnson, R., & Freer, C. (1996). Intelligence and the Frontal Lobe: The Organization of Goal-Directed Behavior. Cognitive Psychology, 30(3), 257–303. https://doi.org/10.1006/cogp.1996.0008

Emrich, S. M., Riggall, A. C., LaRocque, J. J., & Postle, B. R. (2013). Distributed Patterns of Activity in Sensory Cortex Reflect the Precision of Multiple Items Maintained in Visual Short-Term Memory. Journal of Neuroscience, 33(15), 6516–6523. https://doi.org/10.1523/JNEUROSCI.5732-12.2013

Esteban, O., Markiewicz, C. J., Blair, R. W., Moodie, C. A., Isik, A. I., Erramuzpe, A., Kent, J. D., Goncalves, M., DuPre, E., Snyder, M., Oya, H., Ghosh, S. S., Wright, J., Durnez, J., Poldrack, R. A., & Gorgolewski, K. J. (2019). fMRIPrep: A robust preprocessing pipeline for functional MRI. Nature Methods, 16(1), 111–116. https://doi.org/10.1038/s41592-018-0235-4

Frank, M. J., Moustafa, A. A., Haughey, H. M., Curran, T., & Hutchison, K. E. (2007). Genetic triple dissociation reveals multiple roles for dopamine in reinforcement learning. Proceedings of the National Academy of Sciences, 104(41), 16311–16316. https://doi.org/10.1073/pnas.0706111104

Frömer, R., Dean Wolf, C. K., & Shenhav, A. (2019). Goal congruency dominates reward value in accounting for behavioral and neural correlates of value-based decision-making. Nature Communications, 10(1), 4926. https://doi.org/10.1038/s41467-019-12931-x

Garrison, J., Erdeniz, B., & Done, J. (2013). Prediction error in reinforcement learning: A meta-analysis of neuroimaging studies. Neuroscience & Biobehavioral Reviews, 37(7), 1297–1310. https://doi.org/10.1016/j.neubiorev.2013.03.023

Gershman, S. J. (2015). Do learning rates adapt to the distribution of rewards? Psychonomic Bulletin & Review, 22(5), 1320–1327. https://doi.org/10.3758/s13423-014-0790-3

Glasser, M. F., Sotiropoulos, S. N., Wilson, J. A., Coalson, T. S., Fischl, B., Andersson, J. L., Xu, J., Jbabdi, S., Webster, M., Polimeni, J. R., Van Essen, D. C., & Jenkinson, M. (2013). The minimal preprocessing pipelines for the Human Connectome Project. NeuroImage, 80, 105–124. https://doi.org/10.1016/j.neuroimage.2013.04.127

Guo, R., Böhmer, W., Hebart, M., Chien, S., Sommer, T., Obermayer, K., & Gläscher, J. (2016). Interaction of Instrumental and Goal-Directed Learning Modulates Prediction Error Representations in the Ventral Striatum. Journal of Neuroscience, 36(50), 12650–12660. https://doi.org/10.1523/JNEUROSCI.1677-16.2016

Haatveit, B. C., Sundet, K., Hugdahl, K., Ueland, T., Melle, I., & Andreassen, O. A. (2010). The validity of d prime as a working memory index: Results from the “Bergen n-back task.” Journal of Clinical and Experimental Neuropsychology, 32(8), 871–880. https://doi.org/10.1080/13803391003596421

Hamann, S., & Mao, H. (2002). Positive and negative emotional verbal stimuli elicit activity in the left amygdala. NeuroReport, 13(1), 15–19.

Han, S., Huettel, S. A., Raposo, A., Adcock, R. A., & Dobbins, I. G. (2010). Functional Significance of Striatal Responses during Episodic Decisions: Recovery or Goal Attainment? Journal of Neuroscience, 30(13), 4767–4775. https://doi.org/10.1523/JNEUROSCI.3077-09.2010

Howard, J. D., Gottfried, J. A., Tobler, P. N., & Kahnt, T. (2015). Identity-specific coding of future rewards in the human orbitofrontal cortex. Proceedings of the National Academy of Sciences, 112(16), 5195–5200. https://doi.org/10.1073/pnas.1503550112

Izuma, K., Saito, D. N., & Sadato, N. (2008). Processing of Social and Monetary Rewards in the Human Striatum. Neuron, 58(2), 284–294. https://doi.org/10.1016/j.neuron.2008.03.020

Jenkinson, M., Bannister, P., Brady, M., & Smith, S. (2002). Improved Optimization for the Robust and Accurate Linear Registration and Motion Correction of Brain Images. NeuroImage, 17(2), 825–841. https://doi.org/10.1006/nimg.2002.1132

Juechems, K., & Summerfield, C. (2019). Where Does Value Come From? Trends in Cognitive Sciences, 23(10), 836–850. https://doi.org/10.1016/j.tics.2019.07.012

Keramati, M., Dezfouli, A., & Piray, P. (2011). Speed/Accuracy Trade-Off between the Habitual and the Goal-Directed Processes. PLoS Computational Biology, 7(5). https://doi.org/10.1371/journal.pcbi.1002055

Kirchner, W. K. (1958). Age differences in short-term retention of rapidly changing information. Journal of Experimental Psychology, 55(4), 352–358. https://doi.org/10.1037/h0043688

Knutson, B., Fong, G. W., Adams, C. M., Varner, J. L., & Hommer, D. (2001). Dissociation of reward anticipation and outcome with event-related fMRI. NeuroReport, 12(17), 3683–3687.

Kriegeskorte, N., Simmons, W. K., Bellgowan, P. S., & Baker, C. I. (2009). Circular analysis in systems neuroscience – the dangers of double dipping. Nature Neuroscience, 12(5), 535–540. https://doi.org/10.1038/nn.2303

Langdon, A. J., Sharpe, M. J., Schoenbaum, G., & Niv, Y. (2017). Model-based predictions for dopamine. Current Opinion in Neurobiology, 49, 1–7. https://doi.org/10.1016/j.conb.2017.10.006

Leong, Y. C., Radulescu, A., Daniel, R., DeWoskin, V., & Niv, Y. (2017). Dynamic Interaction between Reinforcement Learning and Attention in Multidimensional Environments. Neuron, 93(2), 451–463. https://doi.org/10.1016/j.neuron.2016.12.040

Li, J., Delgado, M. R., & Phelps, E. A. (2011). How instructed knowledge modulates the neural systems of reward learning. Proceedings of the National Academy of Sciences, 108(1), 55–60. https://doi.org/10.1073/pnas.1014938108

Manoach, D. S., Greve, D. N., Lindgren, K. A., & Dale, A. M. (2003). Identifying regional activity associated with temporally separated components of working memory using event-related functional MRI. NeuroImage, 20(3), 1670–1684. https://doi.org/10.1016/j.neuroimage.2003.08.002

McClure, S. M., Berns, G. S., & Montague, P. R. (2003). Temporal Prediction Errors in a Passive Learning Task Activate Human Striatum. Neuron, 38(2), 339–346. https://doi.org/10.1016/S0896-6273(03)00154-5

McClure, S. M., York, M. K., & Montague, P. R. (2004). The neural substrates of reward processing in humans: The modern role of FMRI. The Neuroscientist, 10(3), 260–268. https://doi.org/10.1177/1073858404263526

McDougle, S. D., Butcher, P. A., Parvin, D. E., Mushtaq, F., Niv, Y., Ivry, R. B., & Taylor, J. A. (2019). Neural Signatures of Prediction Errors in a Decision-Making Task Are Modulated by Action Execution Failures. Current Biology, 29(10), 1606–1613.e5. https://doi.org/10.1016/j.cub.2019.04.011

Miller, E. K., & Cohen, J. D. (2001). An Integrative Theory of Prefrontal Cortex Function. Annual Review of Neuroscience, 24(1), 167–202. https://doi.org/10.1146/annurev.neuro.24.1.167

Mumford, J. A., Davis, T., & Poldrack, R. A. (2014). The impact of study design on pattern estimation for single-trial multivariate pattern analysis. NeuroImage, 103, 130–138. https://doi.org/10.1016/j.neuroimage.2014.09.026

Murty, V. P., Shermohammed, M., Smith, D. V., Carter, R. M., Huettel, S. A., & Adcock, R. A. (2014). Resting state networks distinguish human ventral tegmental area from substantia nigra. NeuroImage, 100, 580–589. https://doi.org/10.1016/j.neuroimage.2014.06.047

Palombo, D. J., Hayes, S. M., Reid, A. G., & Verfaellie, M. (2019). Hippocampal contributions to value-based learning: Converging evidence from fMRI and amnesia. Cognitive, Affective, & Behavioral Neuroscience, 19(3), 523–536. https://doi.org/10.3758/s13415-018-00687-8

Pashler, H. (1994). Dual-task interference in simple tasks: Data and theory. Psychological Bulletin, 116(2), 220–244. https://doi.org/10.1037/0033-2909.116.2.220

Pearson, J. M., Heilbronner, S. R., Barack, D. L., Hayden, B. Y., & Platt, M. L. (2011). Posterior cingulate cortex: Adapting behavior to a changing world. Trends in Cognitive Sciences, 15(4), 143–151. https://doi.org/10.1016/j.tics.2011.02.002

Piray, P., Dezfouli, A., Heskes, T., Frank, M. J., & Daw, N. D. (2019). Hierarchical Bayesian inference for concurrent model fitting and comparison for group studies. PLOS Computational Biology, 15(6), e1007043. https://doi.org/10.1371/journal.pcbi.1007043

Power, J. D., Mitra, A., Laumann, T. O., Snyder, A. Z., Schlaggar, B. L., & Petersen, S. E. (2014). Methods to detect, characterize, and remove motion artifact in resting state fMRI. NeuroImage, 84, 320–341. https://doi.org/10.1016/j.neuroimage.2013.08.048

Radulescu, A., Niv, Y., & Ballard, I. (2019). Holistic Reinforcement Learning: The Role of Structure and Attention. Trends in Cognitive Sciences, 23(4), 278–292. https://doi.org/10.1016/j.tics.2019.01.010

Ribas-Fernandes, J. J. F., Solway, A., Diuk, C., McGuire, J. T., Barto, A. G., Niv, Y., & Botvinick, M. M. (2011). A Neural Signature of Hierarchical Reinforcement Learning. Neuron, 71(2), 370–379. https://doi.org/10.1016/j.neuron.2011.05.042

Rissman, J., Gazzaley, A., & D’Esposito, M. (2004). Measuring functional connectivity during distinct stages of a cognitive task. NeuroImage, 23(2), 752–763. https://doi.org/10.1016/j.neuroimage.2004.06.035

Rmus, M., McDougle, S. D., & Collins, A. G. (2021). The role of executive function in shaping reinforcement learning. Current Opinion in Behavioral Sciences, 38, 66–73. https://doi.org/10.1016/j.cobeha.2020.10.003

Satterthwaite, T. D., Ruparel, K., Loughead, J., Elliott, M. A., Gerraty, R. T., Calkins, M. E., Hakonarson, H., Gur, R. C., Gur, R. E., & Wolf, D. H. (2012). Being right is its own reward: Load and performance related ventral striatum activation to correct responses during a working memory task in youth. NeuroImage, 61(3), 723–729. https://doi.org/10.1016/j.neuroimage.2012.03.060

Schuck, N. W., Cai, M. B., Wilson, R. C., & Niv, Y. (2016). Human Orbitofrontal Cortex Represents a Cognitive Map of State Space. Neuron, 91(6), 1402–1412. https://doi.org/10.1016/j.neuron.2016.08.019

Sharpe, M. J., Stalnaker, T., Schuck, N. W., Killcross, S., Schoenbaum, G., & Niv, Y. (2019). An Integrated Model of Action Selection: Distinct Modes of Cortical Control of Striatal Decision Making. Annual Review of Psychology, 70(1), null. https://doi.org/10.1146/annurev-psych-010418-102824

Smittenaar, P., Chase, H. W., Aarts, E., Nusselein, B., Bloem, B. R., & Cools, R. (2012). Decomposing effects of dopaminergic medication in Parkinson’s disease on probabilistic action selection – learning or performance? European Journal of Neuroscience, 35(7), 1144–1151. https://doi.org/10.1111/j.1460-9568.2012.08043.x

Starkweather, C. K., Gershman, S. J., & Uchida, N. (2018). The Medial Prefrontal Cortex Shapes Dopamine Reward Prediction Errors under State Uncertainty. Neuron, 98(3), 616–629.e6. https://doi.org/10.1016/j.neuron.2018.03.036

Sutton, R. S., & Barto, A. G. (1998). Reinforcement learning: An introduction (Vol. 1). MIT Press.

Tustison, N. J., Avants, B. B., Cook, P. A., Zheng, Y., Egan, A., Yushkevich, P. A., & Gee, J. C. (2010). N4ITK: Improved N3 Bias Correction. IEEE Transactions on Medical Imaging, 29(6), 1310–1320. https://doi.org/10.1109/TMI.2010.2046908

Vanderplas, J. M., & Garvin, E. A. (1959). The association value of random shapes. Journal of Experimental Psychology, 57(3), 147–154. https://doi.org/10.1037/h0048723

White, J. K., Bromberg-Martin, E. S., Heilbronner, S. R., Zhang, K., Pai, J., Haber, S. N., & Monosov, I. E. (2019). A neural network for information seeking. Nature Communications, 10(1), 5168. https://doi.org/10.1038/s41467-019-13135-z

Wilson, R. C., Takahashi, Y. K., Schoenbaum, G., & Niv, Y. (2014). Orbitofrontal Cortex as a Cognitive Map of Task Space. Neuron, 81(2), 267–279. https://doi.org/10.1016/j.neuron.2013.11.005

Wolfe, J. B. (1936). Effectiveness of token rewards for chimpanzees. Comparative Psychology Monographs, 12, 72–72.

Yeo, B. T. T., Krienen, F. M., Eickhoff, S. B., Yaakub, S. N., Fox, P. T., Buckner, R. L., Asplund, C. L., & Chee, M. W. L. (2015). Functional Specialization and Flexibility in Human Association Cortex. Cerebral Cortex, 25(10), 3654–3672. https://doi.org/10.1093/cercor/bhu217

